# Cell-specific chromatin landscape of human coronary artery resolves regulatory mechanisms of disease risk

**DOI:** 10.1101/2021.06.07.447388

**Authors:** Adam W. Turner, Sheng’en Hu, Jose Verdezoto Mosquera, Wei Feng Ma, Chani J. Hodonsky, Doris Wong, Gaëlle Auguste, Katia Sol-Church, Emily Farber, Soumya Kundu, Anshul Kundaje, Nicolas G. Lopez, Lijiang Ma, Saikat Kumar B. Ghosh, Suna Onengut-Gumuscu, Euan A. Ashley, Thomas Quertermous, Aloke V. Finn, Nicholas J. Leeper, Jason C. Kovacic, Johan L.M. Björkgren, Chongzhi Zang, Clint L. Miller

## Abstract

Coronary artery disease (CAD) is a complex inflammatory disease involving genetic influences across several cell types. Genome-wide association studies (GWAS) have identified over 170 loci associated with CAD, where the majority of risk variants reside in noncoding DNA sequences impacting *cis*-regulatory elements (CREs). Here, we applied single-cell ATAC-seq to profile 28,316 cells across coronary artery segments from 41 patients with varying stages of CAD, which revealed 14 distinct cellular clusters. We mapped ~320,000 accessible sites across all cells, identified cell type-specific elements, transcription factors, and prioritized functional CAD risk variants via quantitative trait locus and sequence-based predictive modeling. We identified a number of candidate mechanisms for smooth muscle cell transition states and identified putative binding sites for risk variants. We further employed CRE to gene linkage to nominate disease-associated key driver transcription factors such as PRDM16 and TBX2. This single cell atlas provides a critical step towards interpreting *cis*-regulatory mechanisms in the vessel wall across the continuum of CAD risk.

## Introduction

Coronary artery disease (CAD) is the leading cause of death globally and results from injury to the vessel wall and atherosclerotic plaque buildup. Atherosclerotic coronary arteries are complex due to the propensity of multiple cell types to undergo phenotypic switching, including endothelial cells, smooth muscle cells (SMCs), fibroblasts, and various immune cells^1–3^. This has hindered efforts to combat the disease process itself, as currently approved therapies only treat the traditional risk factors. Recent single-cell RNA sequencing (scRNA-seq) analyses have yielded numerous cellular insights into atherosclerosis^4–11^. In particular, lineage-traced scRNA-seq approaches have shown that SMCs transdifferentiate to several distinct phenotypes during atherosclerosis: 1) “fibromyocytes” with fibroblast-like signatures^7^; 2) an intermediate cell state that can become fibrochondrocyte or macrophage-like^9^; or 3) a transitional state giving rise to multiple plaque cell types^10^. Together these studies demonstrate that SMC-derived cells can elicit beneficial or detrimental effects depending on the stage of CAD and/or plaque environment. Despite these advances, the underlying cell-specific regulatory mechanisms remain elusive.

As a complex disease, CAD involves an interplay of environmental and genetic factors over the life course. Genome-wide association studies (GWAS) have now identified over 170 independent CAD loci^12–15^. Many of these are predicted to function in vessel wall processes such as regulation of vascular remodeling, vasomotor tone, and inflammation^16^. The majority of CAD associated single nucleotide polymorphisms (SNPs) reside in non-coding regions and are enriched in *cis*-regulatory elements (CREs)^17^, pointing towards regulatory functions^18^. Since CREs are commonly cell type specific^19,20^, understanding CAD regulatory mechanisms at the cellular level is required to fully interpret the functional impact of risk variants. The Assay for Transpose Accessible Chromatin (ATAC-seq) is a widely adopted approach to systematically detect CREs^21^ and has been conducted in CAD-relevant cultured human coronary artery SMCs^22–24^ and aortic endothelial cells^25^. Thus, single-cell ATAC-seq (scATAC-seq)^26,27^ of human coronary artery samples has the potential to provide a more complete regulatory map to unravel disease mechanisms *in vivo*.

Single-cell epigenomics have been previously applied across various human tissues^8,28–35^ however, to date there are no such reference datasets in coronary artery. In this study, we performed scATAC-based chromatin profiling to uncover ~320,000 candidate CREs in human coronary arteries from 41 patients with varying clinical presentations of CAD. In generating this cell-specific chromatin atlas of the human coronary artery, we deciphered candidate CREs and transcription factors for the major cell types or transition states in the coronary artery. We then applied these profiles to estimate CAD risk variant associations within specific cell types by linking CREs to target gene promoters. Finally, we employed both allele-specific mapping and sequence-based predictive modeling to resolve genetic regulatory mechanisms that could be targeted in the vessel wall.

## Results

### Single-cell ATAC-seq profiling of human coronary artery identifies 14 clusters and captures cellular heterogeneity

We performed scATAC-seq on coronary arteries (left anterior descending artery (LAD), left circumflex artery (LCX), or right coronary artery (RCA)) from 41 patients with various presentations of atherosclerosis using a droplet-based protocol^28^ (**Figure 1a, Supplementary Tables 1,2**). We isolated nuclei from a total of 44 frozen coronary segments using a protocol optimized for frozen tissues^36^ (**Methods**). After sequencing, we performed stringent quality control to retain highly informative cells (**Supplementary Figure 1**). The libraries showed the expected insert size distributions and enrichment of reads at transcription start sites (TSS) (**Supplementary Figure 1**). Aggregating reads from all cells approximated bulk coronary ATAC-seq profiles derived from the same patient, further illustrating the quality of the single-cell dataset (**Supplementary Figure 2**).

**Figure 1.**
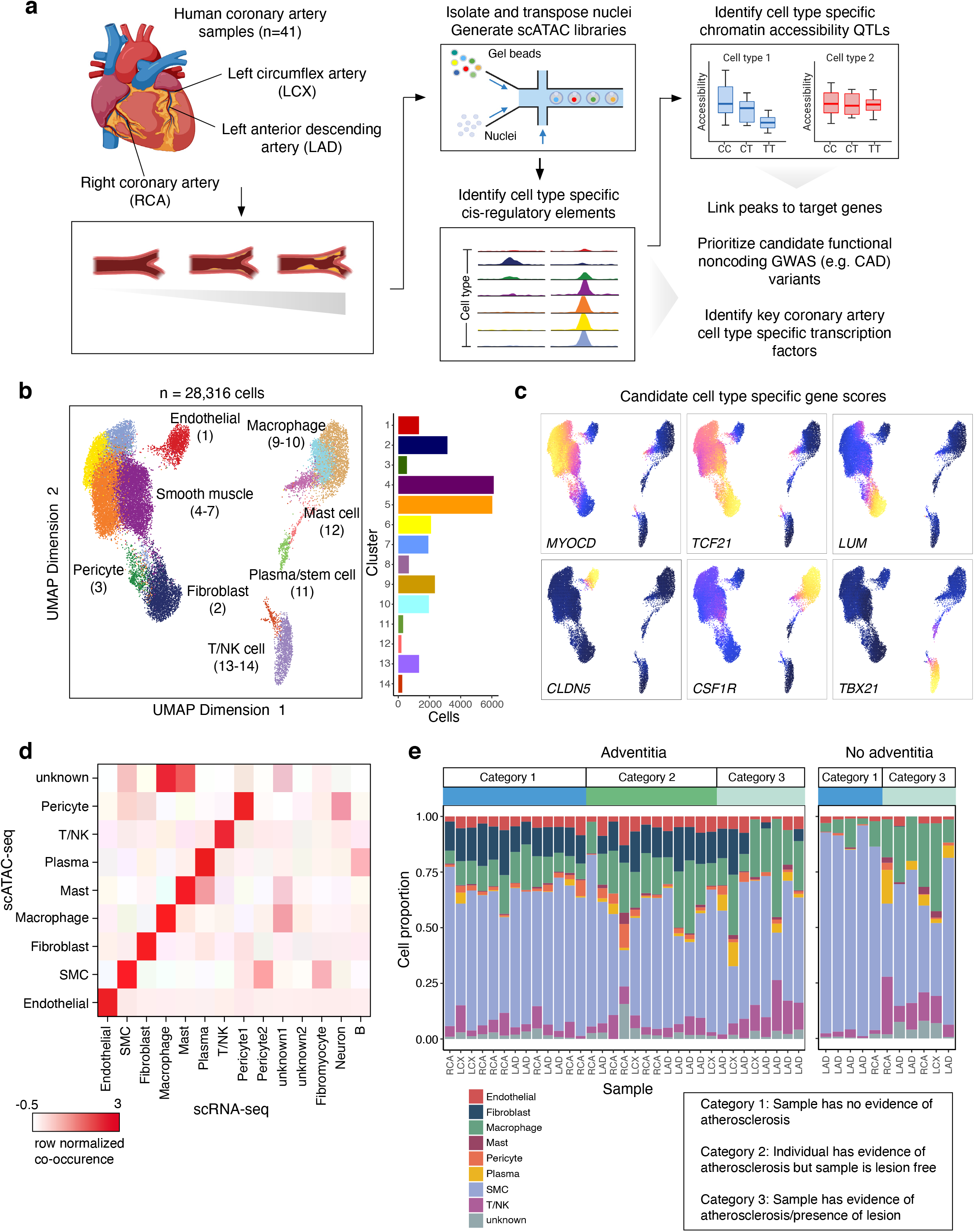
scATAC-seq profiling of 28,316 cells from human coronary arteries reveals cell type chromatin accessibility patterns across 41 individuals. (a) scATAC-seq was performed on nuclei isolated from frozen human coronary artery samples taken from explanted hearts from 41 unique patients. Samples came from segments of either the left anterior descending coronary artery (LAD), left circumflex artery (LCX), or right coronary artery (RCA). After isolation using density gradient centrifugation, nuclei were transposed in bulk and mixed with barcoded gel beads and partitioning oil to generate gel beads in emulsions (GEMs). (b) Uniform manifold approximation and projection (UMAP) and clustering based on single-cell chromatin accessibility identifies 14 distinct coronary artery clusters. Each dot represents an individual cell colored by cluster assignment. (c) UMAP plot of Figure 1b colored by gene score for coronary artery cell type marker genes, including myocardin (*MYOCD*, smooth muscle cells), *TCF21* (smooth muscle cells and fibroblasts), *LUM* (fibroblasts), *CLDN5* (endothelial cells), *CSF1R* (macrophages), and *TBX21* (T cells). (d) Heatmap representing the confusion matrix plot highlighting correspondence between scATAC-seq and scRNA-seq cell type assignments. (e) Distribution of cell types across all of the scATAC-seq samples, divided by whether or not the corresponding sample had an adventitial layer. Schematic in (a) was created using BioRender.

After filtering, we obtained a total of 28,316 cells and identified 14 clusters using iterative Latent Semantic Indexing (LSI) in ArchR^37^ for dimensionality reduction^28,38^ (**Figure 1b**). Importantly, the identified clusters distinguished biological cell types rather than individual donor or other covariates (e.g., age, sex) (**Supplementary Figures 1 and 3**). We assigned each cluster to a coronary artery cell type using gene activity scores, which infer gene expression based on chromatin accessibility at established marker genes^37^. Accessibility at SMC marker genes *MYOCD, MYH11, CNN1, TAGLN*, and *ACTA2* (**Figure 1c, Supplementary Figure 4**) defined four distinct clusters of SMCs, the most abundant cell type in our dataset (57.8 +/− 17.6% of cells, **Figure 1b**). We further identified clusters of endothelial cells (*CLDN5*), fibroblasts (*TCF21, LUM*), macrophages (*CSF1R*), and T cells/natural killer cells (*TBX21*) (**Figure 1c**). Additional cluster annotations included pericytes, plasma (B) cells, and mast cells. Data integration (**Methods**) showed our gene activity score-based annotations were in high agreement with recently reported scRNA-seq annotations from human coronary artery^7^ (**Figure 1d, Supplementary Figure 5**). In general, we observed higher immune cell proportions (cells in clusters 8-14) in atheroma and fibrocalcific coronary artery samples (44.1 +/− 18.8%) relative to non-lesion or healthy controls (17.7 +/− 8.3%), which is consistent with the cellular etiology of atherosclerosis progression^39^ (**Figure 1e, Supplementary Figure 3**). In samples devoid of adventitia (n=11), we observed nearly absent fibroblasts as well as depletion of endothelial cells and pericytes (**Figure 1e**). This is consistent with the expected cell composition of the outer adventitial layer and vasa vasorum^40,41^ and supports the specificity of our cell-type annotations.

### Single-cell ATAC-seq reveals novel cell-type marker genes, regulatory elements and enriched transcription factors

We next applied this coronary artery scATAC-seq dataset to characterize cell type-specific *cis*-regulatory profiles. Using scATAC gene scores we identified 5,121 marker genes across all cell types, which revealed both known and novel cell identity and/or disease response genes (**Figure 2a, Supplementary Table 4**). By aggregating reads from all nuclei we generated a master set of 323,727 peaks, which mostly map to intronic and intergenic sequences as expected (**Supplementary Figure 6** provides peak annotations). Notably, 54% were uniquely accessible in only one or limited cell types (**Figure 2b**), emphasizing the benefits of single-cell profiling to define context-specific regulatory profiles that could be missed in bulk studies.

**Figure 2.**
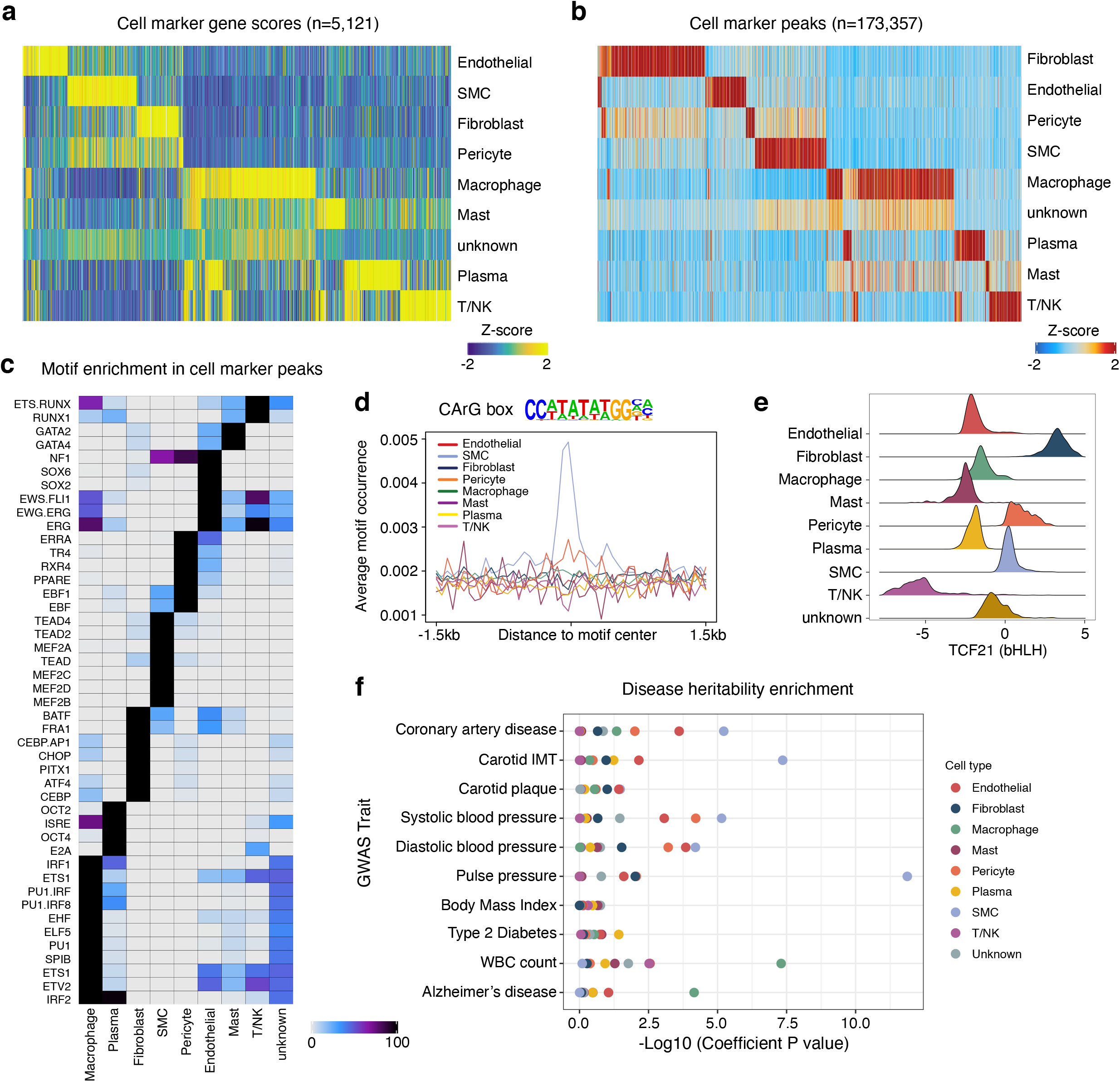
Human coronary artery cell types display distinct gene regulatory processes. (a) Heatmap of coronary cell type marker genes (n=5,121) across each cell type calculated from scATAC-seq gene scores. Each column represents a unique marker gene. The color represents the normalized gene score of the marker genes in cell types. (b) Heatmap reflecting coronary cell type marker peaks that highlight *cis*-regulatory elements specific to only one or very limited cell types. Each column represents an individual marker peak. The color represents the normalized marker peak accessibility in cell types. (c) Heatmap of transcription factor motifs enriched in cell type marker peak sequences. The color represents the normalized motif enrichment score calculated in ArchR using HOMER. (d) Representative motif occurrence plot for the CArG box motif. The CArG box motif, which binds myocardin and serum response factor, is highly enriched in smooth muscle cell accessible chromatin. (e) At the individual cell basis, accessible chromatin is highly enriched for the TCF21 motif in fibroblasts, smooth muscle cells, and pericytes. Transcription factor motif deviations (x-axis) were calculated for each cell using chromVAR. The TCF21 deviations for each cell were integrated based on the cell type (y-axis). (f) LD Score Regression (LDSC) reveals differing enrichment of GWAS SNPs for CAD, hypertension, and non-vascular phenotypes within coronary scATAC cell type peaks.

To investigate the transcription factors (TFs) potentially driving the regulatory profiles/programs in each coronary artery cell type, we performed HOMER^42^ motif enrichment analysis for these marker peaks (**Figure 2c, Supplementary Table 5**). Top enriched motifs in SMCs (MEF2 family^43^, TEAD family^44,45^, CArG box binding myocardin/serum response factor (**Figure 2d**)^46–49^) strongly agree with established SMC TFs in the literature. Similarly, we observed enrichment of ETS and SOX family motifs in endothelial cells^50,51^, PU.1/SPIB and IRF motifs in macrophages^52^, CEBP and AP-1 family members in fibroblasts^53^, RUNX family motifs in T cells^54,55^, and GATA family motifs in mast cells^56^ (**Figure 2c**). Besides defining established cell type specific TFs, we also discovered a number of lesser known coronary TF motifs (e.g., SIX1/2 in fibroblasts and PRDM1 in immune cells). As a complementary approach, we applied chromVAR^57^ on a per-cell level and observed cell-specific motif enrichment (e.g., TCF21 in SMCs and fibroblasts) (**Figure 2e**). Importantly, TCF21 was also highly enriched in fibromyocytes (**Figure 3c**), providing epigenomic-based support for this TF previously shown to drive SMC modulation^7,58^.

**Figure 3.**
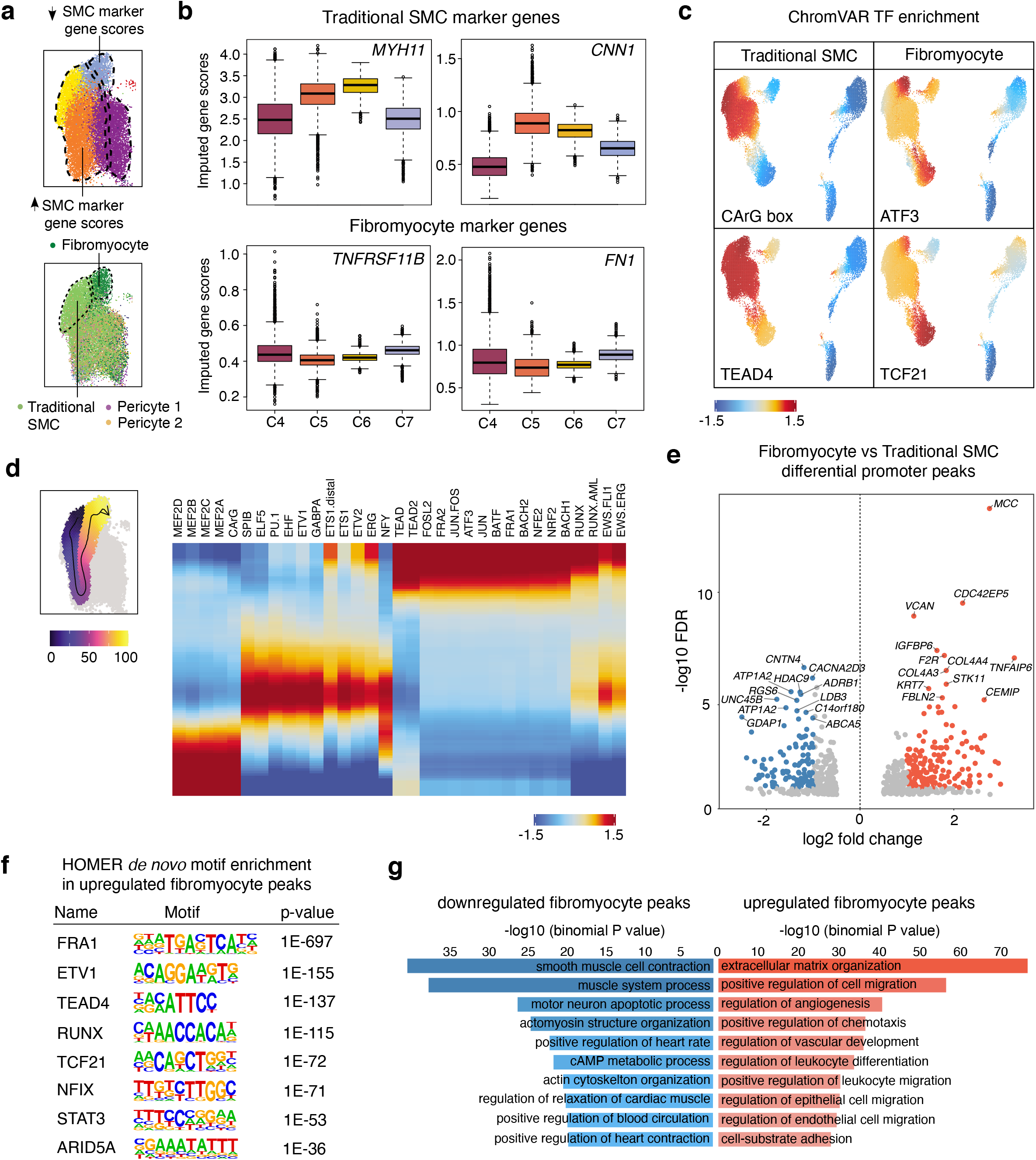
Sub-cluster analysis of smooth muscle cell accessible chromatin identifies fibromyocyte regulatory programs. (a) scATAC UMAP for the 4 SMC clusters (C4-C7). The UMAP was colored by scATAC cluster (top) and by cell type labels assigned by scRNA-seq label transfer (bottom). Integrated scATAC/scRNA UMAP highlights both Fibromyocyte SMCs and traditional SMCs within clusters 4-7. “Pericyte 1” and “Pericyte 2” labels from scRNA-seq were also mixed in clusters 4 and 5. (b) Quantification of imputed scATAC gene scores highlights higher chromatin accessibility at differentiated SMC marker genes *MYH11* and *CNN1* in clusters 5 and 6, and higher accessibility at modulated SMC/Fibromyocyte marker genes *TNFRSF11B* and *FN1* in clusters 4 and 7. (c) ChromVAR transcription factor motif enrichment for differentiated SMC CArG box in traditional SMC and enrichment for ATF3 and TCF21 motifs in modulated SMC/Fibromyocyte and Fibroblast clusters. The TEAD4 motif is enriched in both contractile and modulated SMCs. (d) Left, scatter plot overlay of SMC UMAP depicts the trajectory path from differentiated SMC to modulated SMC/Fibromyocyte sub-clusters (left). Motif enrichment heatmap shows the top enriched motifs across the trajectory pseudo-time (right). Values represent accessibility gene z-scores (e) Volcano plot of differential peak analysis (subset to promoter peaks) comparing Fibromyocyte and traditional SMCs. Fibromyocyte and SMC annotated cells were defined based on RNA label transferring (Methods). Peaks with significant differences were colored light red (Fibromyocyte upregulated) and blue (Fibromyocyte downregulated). (f) Top enriched motifs within the total upregulated Fibromyocyte peaks (5,681) detected using HOMER *de novo* enrichment analysis. (g) Functional annotation of Fibromyocyte upregulated (light red) and downregulated (blue) peaks conducted using GREAT. Top enriched biological processes functional terms are listed.

In order to determine whether cell type-specific accessible regions were enriched for GWAS variants for CAD and other vessel wall phenotypes, we performed cell type Linkage Disequilibrium (LD) Score Regression (LDSC)^59^. CAD and blood pressure GWAS variants were highly enriched in SMC, endothelial cell, and macrophage peaks (**Figure 2f**), while variants for pulse pressure and intimal-medial thickness (cIMT) were specifically enriched in SMC peaks (**Figure 2f**). This is in line with the major contribution of SMC in subclinical atherosclerosis compared to more advanced atherosclerosis involving multiple cell types^3,60^. In contrast, there was limited enrichment for non-vascular traits (**Figure 2f**). Overall, this comprehensive set of coronary artery scATAC-seq chromatin profiles provides a rich landscape to unravel cell-specific regulatory mechanisms in healthy conditions and across diverse diseased stages.

### Characterization of gene regulatory programs in phenotypically modulated SMCs

Given the extensive phenotypic plasticity of SMCs, we next investigated differences in the *cis*-regulatory profiles between contractile and modulated SMCs. While the studies from Alencar et al.^10^ and Pan et al.^9^ both provide compelling evidence of modulated SMC populations, we expanded upon the Wirka et al.^7^ study. This study included human coronary arteries and identified a modulated ‘fibromyocyte population’ (markers *TNFRSF11B* and *FN1*). SMCs in our dataset partition into four sub-clusters (**Figure 1b**), referred to as C4-C7 (**Figure 3a**). Clusters C5 and C6 have greater accessibility in differentiated SMC genes (*MYH11* and *CNN1*), whereas clusters C4 and C7 have greater accessibility in phenotypically modulated SMC marker genes (*TNFRSF11B* and *FN1*) (**Figure 3a-b**). To address potential noise due to sparsity of scATAC-based gene scores, we also derived integrated RNA scores using mutual nearest neighbor integration and label transfer between identified anchors in the scATAC-seq data and Wirka et al.^7^ scRNA-seq data (**Methods, Supplementary Figures 4,5**). This approach provided higher resolution to clearly delineate the SMC-derived fibromyocyte population based on *TNFRSF11B, FN1* and other modulation markers (**Figure 3a, Supplementary Figure 4**). Consistently, chromVAR based TF enrichment revealed highly enriched motifs for AP-1 family members (e.g. ATF3) and TCF21 in the fibromyocyte cluster, which were depleted of differentiated SMC CArG box motif (**Figure 3c, Supplementary Figure 4**). Other TFs such as TEAD4 were enriched in all SMC clusters (**Figure 3c**). Together these results suggest that we can leverage single-cell accessibility profiles to understand regulatory drivers of SMC phenotypic modulation.

In a similar approach, we leveraged chromVAR motif deviations to perform trajectory analyses in SMC clusters. By assigning a path of accessibility from differentiated SMCs towards fibromyocytes (**Figure 3d**), we identified enriched MEF2 and CArG motifs at the start of the trajectory, followed by enrichment of ETS and NFY motifs, then AP-1 and RUNX motifs in fibromyocytes (**Figure 3d**). Using the scRNA-seq integrated data, we further identified 7,802 differentially accessible peaks (5,681 upregulated and 2,121 downregulated) between cells annotated as fibromyocytes vs. traditional/differentiated SMCs (**Methods, Supplementary Tables 6-9**). In particular, we identified 170 significantly upregulated and 108 downregulated promoter peaks in fibromyocytes (**Figure 3e, Supplementary Table 10**). Promoters with higher accessibility in fibromyocytes include several extracellular matrix (ECM) genes (e.g., *VCAN, COL4A3/4* and *TNFAIP6*), previously identified using scRNA-seq^7^. However, we also reveal a number of novel candidate fibromyocyte markers such as the Rho GTPase effector gene, *CDC42EP5*, linked to actin-mediated migration/proliferation^61^. Using HOMER *de novo* motif enrichment, we again observed AP-1 (FRA1), RUNX, TEAD, and TCF21 motifs in upregulated fibromyocyte peaks, but also motifs for inflammatory response factors STAT3 and ARID5A^62^ (**Figure 3f, Supplementary Table 11**). Conversely, MEF2A and CArG box motifs were the top enriched motifs in the downregulated peaks. Genomic region enrichment analysis identified ECM organization and cell migration processes in upregulated fibromyocyte peaks, compared to enrichment for SMC contraction and related processes in downregulated peaks (**Figure 3g**). Together, these scATAC based results confirm the role of TCF21 and also identify new candidate TFs underlying SMC phenotypic modulation during CAD.

### Single cell ATAC provides greater resolution for identifying relevant cell types and targets of CAD GWAS variants

Noncoding GWAS variants are enriched in CREs and often operate in a cell type-specific manner^17,29,63^. We thus prioritized candidate functional CAD GWAS variants^13,15^ using a multitiered approach. To first identify variants in CREs, we overlapped CAD lead variants (and variants in high linkage disequilibrium (r^2^ > 0.8; EUR)) from two recent CAD GWAS metaanalyses^13,15^ with scATAC peaks (**Figure 4a**). This resolved a subset of variants (+/− 50 bp) overlapping both shared and marker peaks (**Figure 4b, Supplementary Tables 12 and 14**), with the majority of CAD SNPs residing within SMC peaks, followed by macrophage, fibroblast, and endothelial cell peaks. Based on these overlaps, we then highlighted target cell types for CAD regulatory variants (**Figure 4c**). Several top candidate CAD variants map to cell typespecific peaks, including rs3918226 at the *NOS3* (endothelial nitric oxide synthase) locus and rs9337951 at the *JCAD* (Junctional cadherin 5 associated)/*KIAA1462* locus^64^ both within endothelial peaks, and rs7500448 at the *CDH13* (T-cadherin) locus within a SMC peak (**Figure 4c**). Other CAD variants map to peaks shared across SMC, fibroblast and pericyte cell types such as rs1537373 at the 9p21 locus (*ANRIL/CDKN2B-AS1*), rs2327429 (upstream of *TCF21*) and rs9515203 at *COL4A2* (**Figure 4c**). The lead variant rs9349379 (*PHACTR1/EDN1*), disrupting a MEF2 binding site^65^, is within a strong SMC and macrophage peak (**Figure 4c**). We also prioritized a number of CAD loci within CREs acting through more than one cell type and confirmed SMC-specificity for previously validated loci such as *LMOD1*^66^ (**Figure 4d**).

**Figure 4.**
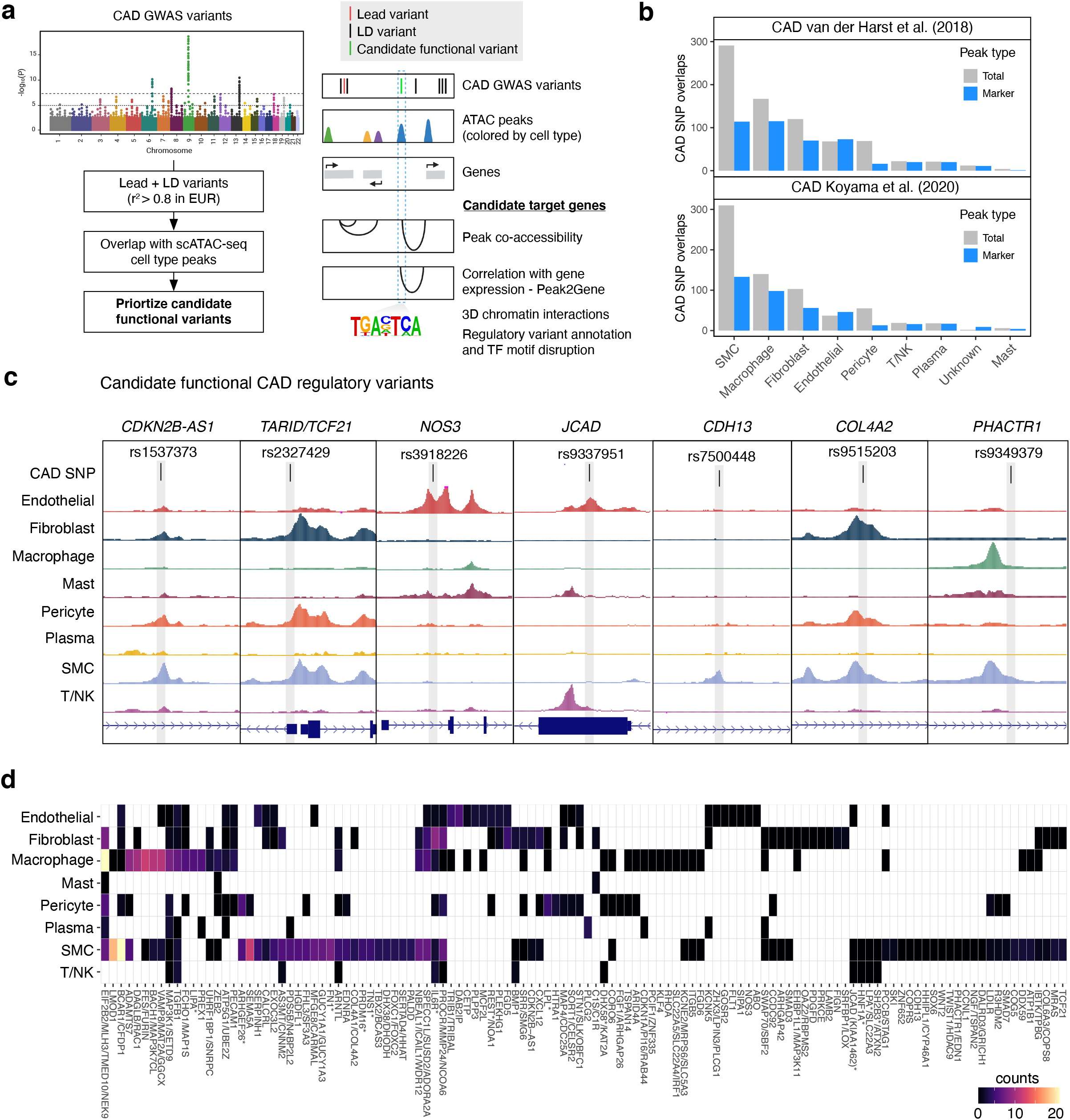
Single-cell chromatin accessibility further resolves mechanisms for functional CAD GWAS loci. (a) To prioritize candidate CAD-associated GWAS variants we used a multitiered strategy, first by taking variants in moderate to high linkage disequilibrium (LD) with the reported lead variants (r^2^ >= 0.8). We next prioritized variants overlapping scATAC peaks and narrowed down the cell type(s) whereby these variants are potentially functioning. Finally we determined whether candidate variants are within transcription factor motifs and linked to target genes through co-accessibility and links to gene expression through scRNA-seq integration (Peak2Gene). (b) Overlap of LD-expanded (r^2^ >=0.8; EUR) CAD GWAS variants (+/− 50 bp) with coronary artery cell type peaks (both from the total peak set and marker peaks). LD-expanded SNPs were obtained from two recent CAD GWAS studies (van der Harst et al. 2018 and Koyama et al. 2020) that performed *trans*-ancestry meta-analysis. (c) Examples of the benefits of scATAC for pinpointing cell types whereby candidate CAD regulatory variants are acting. Highlighted are candidate functional variants at the 9p21 *(CDKN2B-AS1), TARID/TCF21*, *NOS3, JCAD, CDH13, COL4A2*, and *PHACTR1* loci. (d) Heatmap showing number of peaks per cell type overlapping CAD GWAS variants (Only 120 loci shown for clarity). Schematic in (a) was created using BioRender.

Since noncoding variants do not always regulate the nearest gene(s), we also linked candidate variants to target promoters through co-accessibility and scRNA-seq integration (**Methods**, **Figure 4a**). For instance, the SMC peak-containing variant, rs7500448, shows high co-accessibility with the *CDH13* promoter (**Supplementary Figure 7**), which is also an artery-specific eQTL for *CDH13* in GTEx. Another relevant example is rs9985584, located within a strong fibroblast peak 3’ of *VEGFA*, which is highly co-accessible with the *VEGFA* promoter (**Supplementary Figure 7**). In an orthogonal approach, we combined co-accessibility and RNA integration to identify peaks where accessibility correlates with target gene expression (Peak2Gene links). We identified a total of 148,617 Peak2Gene links when aggregating all cell types (**Supplementary Figure 7**), including for many CAD risk variants (**Supplementary Tables 13, 15**). Together, these single-cell chromatin annotations refine candidate regulatory mechanisms at CAD GWAS loci, which can be functionally validated in the appropriate cell types.

### Prioritizing candidate functional variants within coronary artery cell types

Chromatin accessibility quantitative trait locus (caQTL) mapping is a powerful association analysis to resolve candidate GWAS regulatory mechanisms^67–72^. We thus calculated caQTLs in our dataset in four major coronary cell types (SMC, macrophages, fibroblasts, and endothelial cells). Given our modest sample size (n=41), we used RASQUAL^73^ for caQTL mapping to capture both population and allele-specific effects (**Methods**), as done previously for cultured coronary artery SMCs^74^. As expected, the number of QTLs discovered per coronary cell type was proportional to the respective number of annotated cells (**Figure 5a and Supplementary Figure 8**), with the most belonging to SMCs (2,070 at 5% FDR). To determine whether these caQTLs regulate gene expression, we queried these variants for eQTL signals in GTEx artery tissues (coronary, aorta, and tibial). Out of the 2,040 unique SMC caQTLs (5% FDR), 47% were significant eQTLs (GTEx 5% FDR) in at least one GTEx arterial tissue. Most of the coronary SMC caQTLs that are GTEx eQTLs are shared across all artery types (**Figure 5b**). We also identified 71% concordant coronary artery caQTL and eQTL effect sizes (**Figure 5c**), which is consistent with reported findings in human T cells^69^.

**Figure 5.**
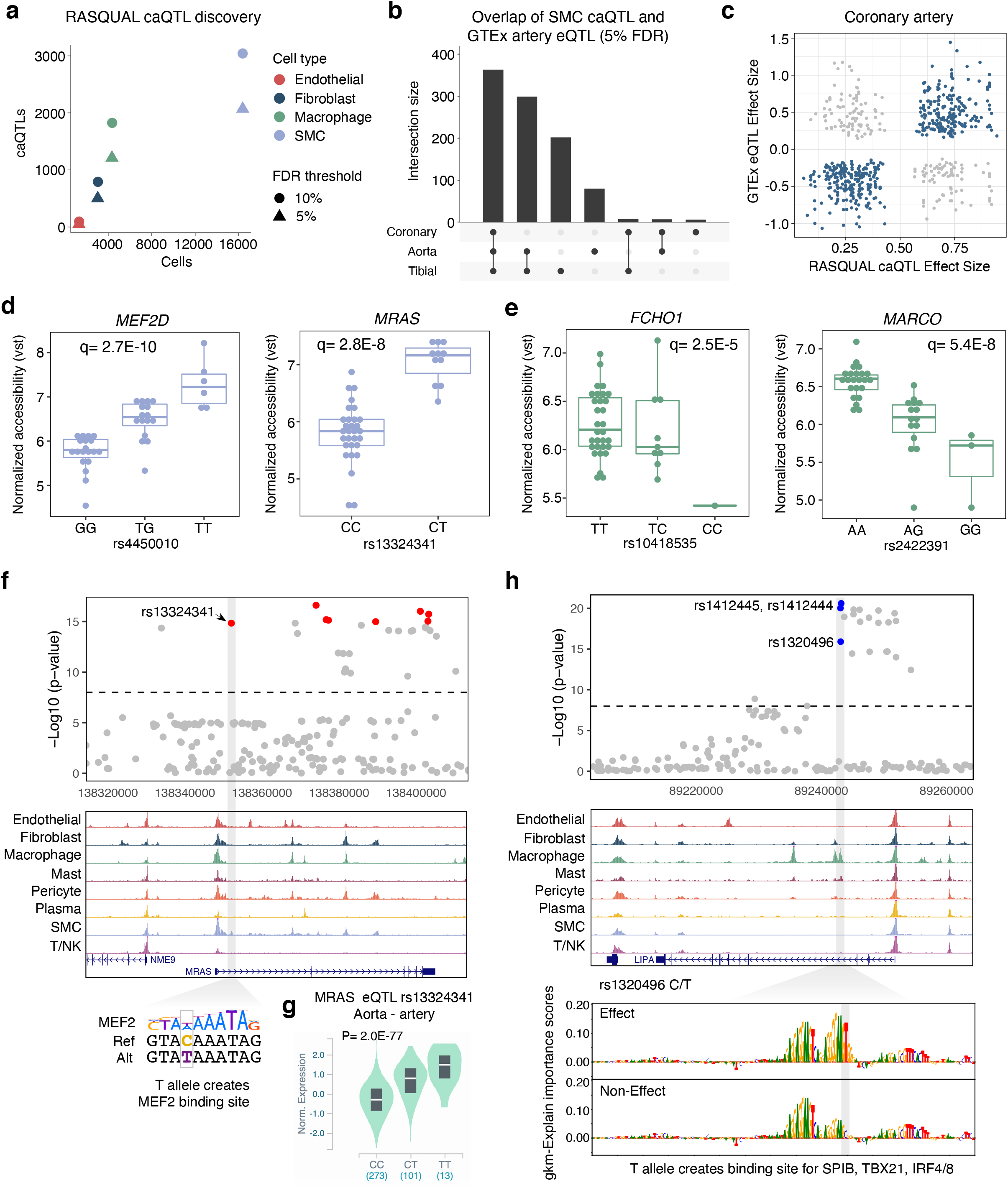
Identification of genetic variants that regulate chromatin accessibility within coronary artery cell types. (a) The number of chromatin accessibility quantitative trait loci (caQTLs) identified using RASQUAL (at both 5% and 10% FDR cutoffs) within a cell type is proportional to the number of annotated cells. The color represents the cell type and shape represents the FDR cutoff. (b) UpSet plot for smooth muscle cells caQTLs that are expression quantitative trait loci in GTEx arterial tissues. Bars represent the intersection size for overlap of eQTLs between coronary artery, aorta, and tibial artery. (c) Comparison of RASQUAL effect sizes with GTEx effect sizes (beta). (d) Boxplots highlighting smooth muscle cell normalized accessibility for *MEF2D* and *MRAS* caQTL variants and (e) macrophage normalized accessibility for *FCHO1* and *MARCO* caQTL variants. The q-values represent the lead SNP q-value that was output from RASQUAL for the respective peak. (f) Example genome browser tracks showing CAD-associated caQTL at the *MRAS* locus in a smooth muscle cell specific peak. The T allele for rs13324331 creates a MEF2 putative binding site. (g) In GTEx artery (aorta shown here) the T allele for rs13324331 is highly associated with increased *MRAS* mRNA levels. cis-eQTL p-value shown. (h) Example of prioritization of functional CAD variants using lsgkm machine learning based prediction. The rs13202496 variant at the *LIPA* locus (chromosome 10) resides in a strong macrophage peak. The T allele is predicted to create a putative SPIB binding site and increased chromatin accessibility. Feature importance score tracks for effect and non-effect alleles are visualized by gkmExplain (Methods).

We next applied these cell type caQTLs to further dissect CAD related mechanisms in the vessel wall. One example is rs4450010 at the *MEF2D* migraine/cardiovascular-associated gene within a SMC peak. The rs4450010-T allele creates a TEF1 (TEAD) binding site and correlates with both increased peak accessibility (**Figure 5d**) and increased *MEF2D* RNA expression in GTEx arterial tissues (**Supplementary Figure 8**). Several CAD GWAS variants were significant caQTLs in SMC or macrophages. For example, rs13324341 within intron 1 of *MRAS* (muscle RAS oncogene homolog, NS13), also in a DNase site^75^, is both a SMC caQTL and strong eQTL in GTEx arterial tissues (**Supplementary Figure 7**). The rs13324341 minor allele T (increased CAD risk) creates a MEF2 binding site (**Figure 5e**) and correlates with both increased accessibility (**Figure 5d**) and increased *MRAS* mRNA levels (**Figure 5g**). Other top CAD GWAS-overlapping caQTLs include, among others, rs73551705 (*BMP1*) and rs17293632 (*SMAD3*,) in SMCs and rs72844419 (*GGCX*) and rs10418535 (*FCHO1*) in macrophages (**Supplementary Figure 8 and Figure 5e**).

To complement our QTL-based approach, we employed a machine learning-based strategy to assign sequence importance scores to CAD variants (10,117 tested) with effects on chromatin accessibility^76^. Across three similar approaches (GkmExplain^77^, gkmpredict, deltaSVM^78^) we identified 127 high- or moderate-confidence CAD variants with predicted functional effects on chromatin accessibility (**Supplementary Table 21**). 102 (80%) had functional probability scores >0.6 in RegulomeDb 2.0 and were annotated by enhancer, promoter, and TF ChIP-seq enrichment as well as motif disruption (**Supplementary Table 22**). About half of these variants were predicted to be functional in a single cell-type. One representative CAD variant, rs1320496 (*LIPA*), resides in a strong macrophage-specific peak, with the T allele (increased CAD risk) creating putative binding sites for SPIB, TBX21, and IRF4/8 (**Figure 5h**). Another intergenic SNP, rs10418535-C/T (between *MAP1S* and *FCHO1*), resides in a macrophage-specific peak with Peak2Gene links to *FCHO1*. The rs10418535-C allele (increased CAD risk) disrupts a PU.1/IRF motif and is predicted to attenuate chromatin accessibility (**Supplementary Figure 9**). rs10418535 is also a macrophage caQTL with a positive effect for the T allele, consistent with the deltaSVM prediction (**Figure 5e, Supplementary Figure 9**). Together, we demonstrate celltype caQTL mapping and machine learning are complementary approaches to pinpoint candidate functional disease risk variants at high resolution.

### Epigenomic profiles highlight PRDM16 and TBX2 as CAD transcription factors

Epigenomic profiles in disease-relevant tissues have been shown to resolve the correct target gene(s) at GWAS risk loci, which are often incorrectly annotated to the nearest gene^79^. Importantly, our data nominates two previously unannotated TFs as candidate causal genes at their respective CAD loci. The first locus on chr17 harbors dozens of tightly linked CAD variants within peaks in the *BCAS3* gene (**Figure 6a**). However, Peak2Gene analysis demonstrates stronger links between these peaks and TBX2 expression in coronary arteries. We also observed colocalization of CAD variants with *TBX2* cis-eQTLs in the Stockholm-Tartu Atherosclerosis Reverse Network Engineering Task (STARNET) aortic root tissue (**Supplementary Figure 10**). At the second locus on chr1, several linked CAD variants are located within SMC peaks 5’ of the *ACTRT2* gene and are highly correlated with *PRDM16* and *LINC00982 (PRDM16* divergent transcript) expression, but not other genes at the locus (**Figure 6b**). This locus also harbors an independent missense CAD-associated SNP (rs2493292; p.Pro634Leu) in exon 9 of *PRDM16*, suggesting both noncoding and coding effects on *PRDM16* expression (**Figure 6b**).

**Figure 6.**
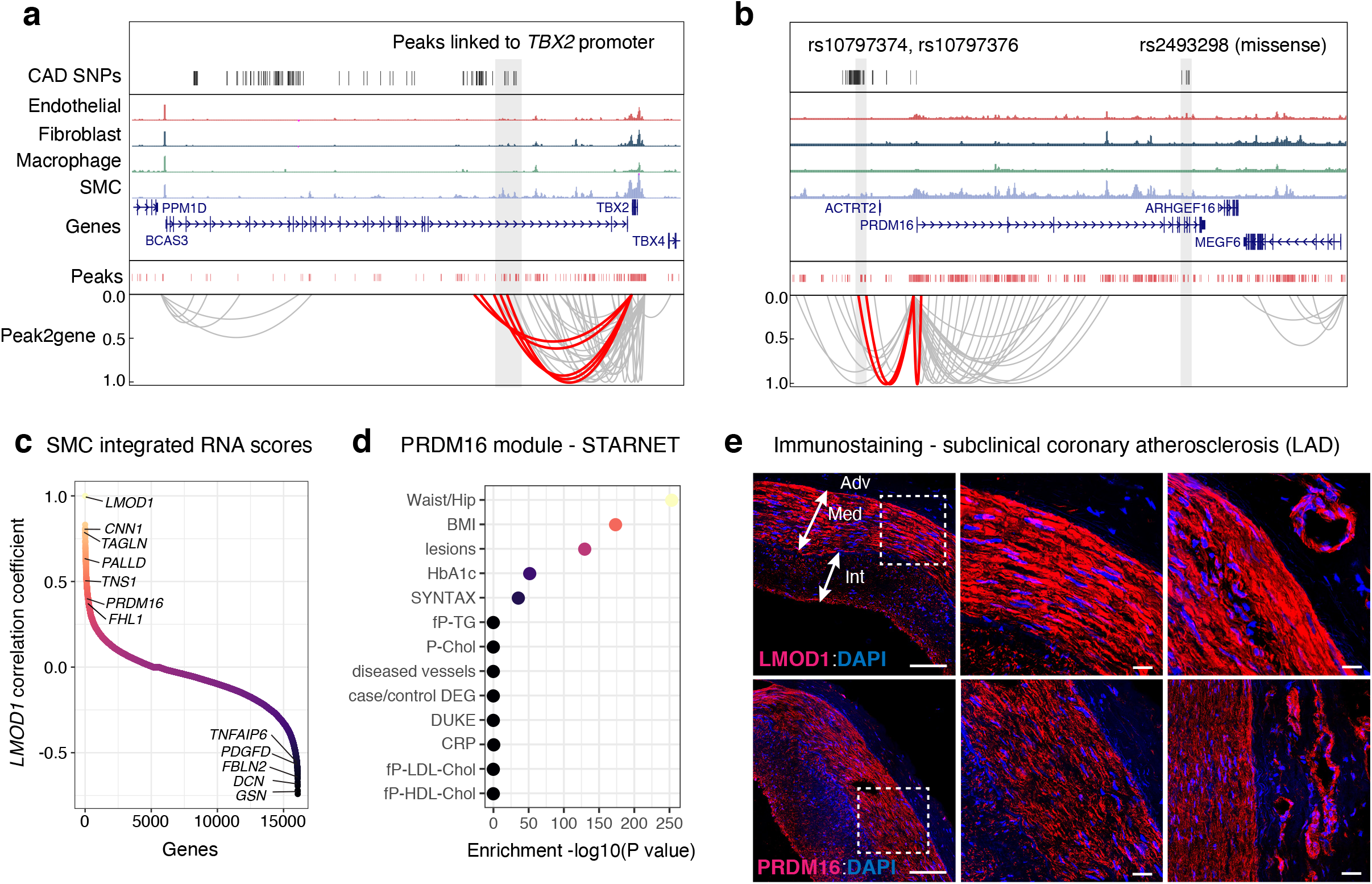
PRDM16 and TBX2 are CAD-associated key driver transcriptional regulators in SMCs. (a-b) Genome browser tracks highlighting the association between CAD associated SNPs and SMC marker genes (a, *TBX2*; b, *PRDM16*) through co-accessibility (peak2gene) detected by scATAC-seq data (Methods). The red loops represent the association between *TBX2* or *PRDM16* promoter and CAD associated SNPs. (c) Correlation coefficients of scATAC/scRNA integration scores gene expression levels between *LMOD1* and genome-wide coding genes in SMCs. Genes were ranked by the correlation coefficient with *LMOD1*. Representative positive and negative correlated SMC genes are labeled. (d) Clinical trait enrichment for PRDM16 containing module in subclinical mammary artery in STARNET gene regulatory network datasets. Pearson ‘s correlation p-values were aggregated for each coexpression module using Fisher’s method. (e) PRDM16 and LMOD1 immunostaining (red) of medial SMC in subclinical coronary artery atherosclerosis samples from left anterior descending (LAD). Middle panels represent higher magnification of highlighted boxes. Right panels depict positive arteriole staining in vasa vasorum. DAPI marks nuclei. Scale = 100 um. Adv: adventitia, Med: media, Int: intima.

Both PRDM16 and TBX2 are scATAC SMC marker genes and remarkably *PRDM16* is one of the top SMC marker genes along with known SMC gene *LMOD1* (**Supplementary Figure 4, Supplementary Table 4**). Given the similar gene score enrichment of *PRDM16* and *LMOD1* in SMC, we ranked *PRDM16* by correlating all SMC gene scores and integrated RNA scores with *LMOD1* (**Figure 6c**). Interestingly, *PRDM16* was modestly correlated with traditional SMC markers, and negatively correlated with fibromyocyte marker genes. This may implicate *PRDM16* as a SMC injury-response gene as opposed to a SMC identity marker gene. To gain further insight into these two TFs, we queried STARNET gene regulatory networks across 7 cardiometabolic tissues (n=600), which revealed both PRDM16 and TBX2 as significant key drivers in artery tissues (**Supplementary Table 26, Supplementary Figure 10**). In subclinical artery, the PRDM16-regulated module was highly enriched for the presence of atherosclerotic lesions and CAD severity, as well as metabolic clinical traits (**Figure 6d, Supplementary Figure 10**). Finally, we confirmed PRDM16 expression via immunofluorescence of human subclinical atherosclerotic coronary artery, with LMOD1 as a positive control. Similar to LMOD1, PRDM16 is primarily located in the medial coronary layer and also expressed in small vessels in the vasa vasorum (**Figure 6c, Supplementary Figure 10**). While we highlight these two examples, this coronary dataset can be similarly utilized to prioritize mechanisms at many other CAD loci.

## Discussion

In this study we have generated the first single-cell atlas of human coronary artery chromatin accessibility for over 40 patients encompassing healthy and atherosclerotic samples, which captures gene regulation *in vivo*. Over half of the 323,767 identified *cis*-regulatory elements (54%) are unique to a specific cell type or limited number of cell types, underscoring the power of single-cell epigenomics for resolving unique cell type regulatory processes. Our scATAC results also provide novel insights into SMC phenotypic modulation. More specifically, we discovered accessible regions, genes and putative TF motifs that may drive the transition of native SMCs towards modulated SMCs (e.g. fibromyocytes). Finally, using an integrative statistical genetics and machine learning approach we prioritized cell-specific candidate regulatory variants and mechanisms underlying CAD loci.

There are now over 170 genetic loci associated with CAD risk, primarily located within noncoding genomic regions^13,15^. This single-cell coronary artery epigenomic landscape provides a valuable resource to disentangle the target cell type(s), candidate causal genes and variants at the expanding number of CAD risk loci in diverse populations. For example we highlight a top caQTL rs13324341 in the *MRAS* locus which alters a MEF2 binding site in SMCs. Given the role of *MRAS* in Noonan syndrome-associated cardiomyocyte hypertrophy^80^, these results may provide clues into SMC growth responses during CAD. We also highlight top predicted CAD regulatory variants acting in one or more cell types (e.g. rs1320496 at *LIPA* in macrophages). This dataset can also be leveraged to interrogate GWAS loci for related common vascular diseases (e.g., hypertension or coronary artery calcification).

By taking into account co-accessibility and scRNA-seq integration, we systematically link CAD risk variants to target gene promoters. This is critical given that GWAS variants are estimated to only target the nearest gene ~50% of the time. For example, using this approach we nominate two TFs-PRDM16 and TBX2-as top candidate genes at their respective loci. *PRDM16* is a top SMC marker gene from scATAC gene scores, however it may not be limited to marking SMC identity. PRDM16 (MEL1) is a TF known for roles in metabolism and controlling brown fat-to-skeletal muscle switches^81–83^. However, *PRDM16* is enriched in GTEx arterial tissues and was identified as a key driver gene in STARNET artery tissue, consistent with our scATAC data. PRDM16 regulates TGF-beta signaling^84^ through direct interactions with Smad^85^ and SKI^86^ proteins, both of which are associated with CAD^13^. PRDM16 may play key roles in endothelial cells in arterial flow recovery^87^. Similarly, *TBX2* is enriched in GTEx arterial tissues and SMC clusters in our dataset, consistent with prior studies showing that Tbx2 activates SRF^88^. TBX2 also links to relevant CAD pathways such as BMP, TGF-beta, and FGF signaling^89^. Functional follow-up studies to investigate target binding sites and affected SMC processes for these TFs may reveal new mechanisms of disease risk.

While this study provides high-resolution insights into coronary artery gene regulatory signatures using valuable human tissue samples, there are some known limitations. Given the lack of available lineage-tracing scATAC datasets, we cannot fully annotate intermediate cell types or precisely resolve their origins and fates during atherosclerosis^9,15^ For example, some SMC-derived cells may be incorrectly annotated in T cell clusters, consistent with Alencar et al.^10^ and Hansson et al.^90^. Also, it is worth noting that we captured more cells from subclinical lesions compared to advanced atherosclerotic lesions, which potentially reflects higher difficulty in nuclei extraction for diseased samples. This disparity may underrepresent distinct classes of T cells and NK cells, known to play fundamental roles in atherosclerosis^4,5^ Finally, given our modest sample size for QTL based studies, we were underpowered to discover a large number of caQTLs for less abundant cell types or transition states. Decreasing costs and adoption of novel single-cell and spatial sequencing technologies may further improve discovery of regulatory variants and mechanisms through multi-modal and integrative approaches^91,92^.

In summary, we provide an atlas of chromatin accessibility in both healthy and atherosclerotic human coronary arteries. These cell type-specific epigenomic profiles now illuminate *cis-* regulatory programs at basepair resolution to further our understanding of cell plasticity and heritable disease risk in the coronary vessel wall. We anticipate this will provide a valuable resource to functionally interrogate causal disease processes and inform pre-clinical studies to circumvent atherosclerosis.

## Online Methods

### Coronary artery tissues and human subjects

Freshly explanted hearts from orthotopic heart transplantation recipients were procured at Stanford University under approved Institutional Review Board protocols and written informed consent. Hearts were arrested in cardioplegic solution and rapidly transported from the operating room to the adjacent lab on ice. The proximal 5-6 cm of three major coronary vessels (left anterior descending (LAD), left circumflex (LCX), and right coronary artery (RCA)) were dissected from the epicardium on ice, trimmed of surrounding adipose (and in some samples the adventitia), rinsed in cold phosphate buffered saline, and rapidly snap frozen in liquid nitrogen. Coronary artery samples were also obtained at Stanford University (from Donor Network West and California Transplant Donor Network) from non-diseased donor hearts rejected for heart transplantation and procured for research studies. Hearts were arrested in cardioplegic solution and transported on ice following the same protocol as hearts used for transplant. Explanted hearts were generally classified as ischemic or non-ischemic cardiomyopathy and prior ischemic events and evidence of atherosclerosis was obtained through retrospective review of electronic health records at Stanford Hospital and Clinics. The disease status of coronary segments from donor and explanted hearts was also evaluated by gross inspection at the time of harvest (for presence of lesions), as well as histological analysis of adjacent frozen tissues embedded in OCT blocks. Frozen tissues were transferred to the University of Virginia through a material transfer agreement and Institutional Review Board approved protocols. All samples were then stored at −80°C until day-of-processing.

### Isolation of nuclei for single-cell ATAC-seq

We performed single-cell ATAC on four coronary artery samples per day. For nuclear isolation we used a similar protocol to Omni-ATAC^36^ that was optimized for frozen tissues and reported lower mitochondrial reads. Approximately 50 mg of frozen human coronary artery tissue was broken into small pieces/powder using chilled Mortar and Pestle on dry ice and liquid nitrogen. This was transferred back to a chilled 1.5 mL microcentrifuge tube and kept on dry ice until all samples were broken down. We added 1 mL of cold 1X Homogenization Buffer (5 mM CaCl_2_, 3 mM Mg(Ac)2, 10 mM Tris pH 7.8, 320 mM sucrose, 0.1 mM EDTA, 0.1% NP-40, 0.1 mg/mL BSA, Roche cOmplete protease inhibitor) and the tubes were gently inverted 5 times and the powder gently pipetted up and down using a wide-bore 1 mL pipette tip set to a volume of 1 mL. The samples were then immediately transferred to cold, pre-chilled 1 mL glass dounce homogenizers on ice.

We performed Dounce homogenization (10 strokes with Pestle A (Loose) and 20 strokes with Pestle B (Tight)) on ice and passed the lysate through a 70 μm Falcon strainer (Corning). The flow-through was collected and transferred to a chilled 2 mL Lo-Bind microcentrifuge tube (Eppendorf) and centrifuged at 4°C for 1 minute at 100 g. This supernatant was transferred to a new 2 mL Lo-Bind microcentrifuge tube (Eppendorf) and an OptiPrep/sucrose (Sigma) gradient was generated the same way as in the Omni-ATAC protocol^36^. 400 μl of sample was first mixed thoroughly with 400 μl of 50% Iodixanol by pipetting. Next, we layered 600 μl of 29% Ioxinadol underneath and then 600 μl of 35% Iodixanol underneath the 29% layer. Samples were centrifuged for 20 minutes at 10,000 g at 4°C and the top layers were aspirated down to within 300 μl of the nuclei band.

We then carefully took the band containing the nuclei (setting the pipette volume to 100 μl) and added the nuclei to 1.3 mL of cold Nuclei Wash Buffer (10 mM T ris-HCl (pH 7.4), 10 mM NaCl, 3 mM MgCl_2_, 1% BSA, 0.1% Tween-20) in a 1.5 mL Lo-Bind microcentrifuge tube. The microcentrifuge tube was inverted gently 5 times, nuclei gently mixed by pipetting (setting the pipette volume to 1 mL), and contents passed through a 40 μm Falcon cell strainer (Corning) into a new 1.5 mL Lo-Bind microcentrifuge tube (Eppendorf). Nuclei were pelleted by centrifugation for 5 minutes at 500 g at 4°C and supernatant carefully removed. Finally, this nuclei pellet was gently resuspended in 100 μl of the Nuclei Buffer provided with the kit (diluted from 20X Stock to 1X working concentration with nuclease-free water) by gently pipetting up and down. Samples and nuclei were kept on ice for all steps of the nuclear isolation. For each sample we measured the nuclei concentration by taking the mean of two separate counts using Trypan blue (Thermo Fisher) and the Countess II instrument (Thermo Fisher). Post cell lysis we generally observed less than 5% Live cells when visualizing with the Countess, consistent with proper lysis.

### Single-cell ATAC reaction setup

For the first four samples processed we used the 10x Genomics Chromium Single Cell ATAC Library & Gel Bead Kit for four reactions (PN-1000111). For all subsequent experiments we used the 10x Genomics Chromium Next GEM Single Cell ATAC Reagent Kits, v1.1 (PN-1000175 and PN-1000176). For each single-cell ATAC transposition reaction we mixed 7 μl of ATAC Buffer B, 3 μl of ATAC Enzyme, and 5 μl of nuclei (in Diluted Nuclei Buffer). We then set the pipette volume to 10 μl and mixed by pipetting up and down six times. Transposition was performed in a PCR machine for 60 minutes at 37°C in a 15 μl reaction volume (50°C lid temperature).

### Single-cell ATAC GEM preparation

The full protocols for the single cell ATAC-seq data generation are available at the following link: https://support.10xgenomics.com/single-cell-atac.

We first performed a pilot study (4 samples) using the original Chromium Single Cell ATAC Reagent Kit protocol (10x Genomics, PN-1000111). The remaining samples were processed using the Chromium Next GEM Single Cell ATAC Reagent Kit v1.1 (10x Genomics, PN-1000175 and PN-1000176) protocol. Transposed nuclei (15 μl) were mixed with Barcoding Enzyme/Reagent Master Mix (60 μl) for a total of 75 μl per sample. This Transposed Nuclei + Master Mix volume was subsequently added to the 10x Genomics Chromium GEM chip. 50 μl of Gel Beads and 40 μl of Partitioning Oil were subsequently added to the chip and the chip covered with the 10x Gasket. The assembled chip was then run on the 10x Genomics Chromium Controller instrument to generate Gel Beads in Emulsions (GEMs). 100 μl of GEMs were then incubated in a thermal cycler (105°C lid temperature) using the following cycling conditions: 72°C for 5 minutes; 98°C for 30 seconds; 12 cycles of 98°C for 10 seconds, 59°C for 30 seconds, 72°C for 1 minute; then holding at 15°C. After incubation, GEMs were cleaned up using Dynabeads MyOne and SPRIselect reagent to produce a volume of 40 μl.

### Single-cell ATAC library generation

Each sample was uniquely indexed with the Chromium i7 Multiplex Kit N, Set A, 96 rxns (10x Genomics) PN-1000084. The index sequences for this kit are listed in **Supplementary Table 3**. The sample index PCR reaction was prepared by mixing 40 μl of single-cell ATAC library with 57.5 ul of Sample Index PCR Mix and 2.5 μl of an individual primer from the Chromium i7 Sample Index N, Set A kit. The subsequent PCR amplification conditions were as follows: 98°C for 45 seconds; then 11 cycles of 98°C for 20 seconds, 67°C for 30 seconds, and 72°C for 29 seconds; 72°C for 1 minute; then holding at 4°C. After the sample index PCR we performed a final double sided size selection using SPRIselect beads. Before sequencing all single-cell ATAC libraries were run on the Agilent TapeStation High Sensitivity D1000 or D5000 ScreenTape. Library sizes are shown in **Supplementary Figure 1**.

### Single-cell ATAC library sequencing

Single-cell ATAC libraries were shipped on dry ice to the Genome Core Facility at the Icahn School of Medicine at Mt. Sinai (New York, New York) for sequencing on an Illumina NovaSeq 6000. 40 libraries were sequenced using a NovaSeq S1 flow cell (100 cycles, 2 x 50 bp) and 4 libraries were sequenced using a NovaSeq S Prime (SP) flow cell (100 cycles, 2 x 50 bp).

### Statistical analyses

The statistical tests performed are listed in the respective figure legends or sections of the Methods.

### Genome annotations

All sequence alignments and annotations are with respect to the hg38 human reference genome.

### Raw scATAC data processing and quality control

In-house single-cell ATAC-seq (scATAC) data was pre-processed using the 10x Genomics pipeline (Cell Ranger ATAC version 1.2.0^37^) using the hg38 genome and default parameters. Samples from different patients were pre-processed separately. Individual cells with high quality were kept for downstream analysis (TSS enrichment >=7, unique barcode number >= 10,000 and doublet ratio < 1.5). QC measurements and filtering were conducted using ArchR (v1.0.1)^37^.

### Clustering of coronary artery scATAC data

scATAC reads from different individuals were combined at the single cell level and then mapped to each 500 bp bin across the hg38 reference genome for dimensionality reduction and clustering. The dimensional reduction was conducted using a latent semantic indexing (LSI) algorithm and top 25,000 bins with highest signal variance across individual cells were selected as input. The top 30 dimensions were selected for cell clustering. Cell clusters were identified by a shared nearest neighbor (SNN) modularity optimization-based clustering algorithm from the Seurat^93^ package). Batch effect removal was conducted using the Harmony (v1.0)^94^ package. We did not observe any improvement after batch correction using Harmony^94^

### Single-cell ATAC gene scores and cell-type specific genes

The chromatin accessibility within a gene body as well as proximally and distally from the TSS was used to infer gene expression via computation of a “Gene Score” using the default method in ArchR. The gene score profiles for all cells were subsequently used to generate a gene score matrix. The gene score matrix was also integrated with single-cell RNA-seq (scRNA) expression data (described below). Finally, a cell type annotation for each cluster was assigned using gene scores for cell type marker genes and later validated or further refined through scRNA-seq label transfer (**Figure 1b-d**).

Cell-type specific marker genes in our scATAC data (genes with significantly higher chromatin accessibility in a cluster than in other clusters) were identified using Wilcoxon rank-sum test and the genes with (Benjamini-Hochberg) adjusted p-value <= 0.01 and fold change >=2 were selected. The z-normalized gene scores for the cell-type specific genes were plotted as heatmap (**Figure 2a**).

### Single-cell RNA-seq processing and integrative analysis with scATAC

We integrated the coronary scATAC-seq dataset (28,316 cells) with previously published human coronary artery single-cell RNA-seq (scRNA-seq) from Wirka et al.^7^. The preprocessed scRNA-seq data was downloaded from Gene Expression Omnibus (GEO) and processed using Seurat^93^ as described in the study. Genes expressed in less than 5 cells were filtered out. Cells with <= 500 or >= 3500 genes were also trimmed from the dataset as they likely represent defective cells or doublet/multiplet events. Moreover, cells containing >= 7.5% of reads mapping to the mitochondrial genome were discarded as low quality/dying cells often exhibit high levels of mitochondrial contamination. Upon discarding poor quality cells, 11756 high quality cells remained for further analysis. Read counts were normalized using Seurat’s global-scaling method that normalizes gene expression measurements for each cell by the total expression, multiplies them by a 10000-scaling factor and log-transforms them. Upon finding the 2000 most variable genes in the data, dimensionality reduction was performed using PCA. The top 10 PCs were further used for UMAP visualization and cell clustering (using a shared nearest neighbor (SNN) modularity optimization-based algorithm in the Seurat package). The cluster specific genes (marker genes) for each cluster were identified with Seurat default method. The cell types of clusters were assigned according to the comparison between the cluster specific genes and the cell-type specific gene lists provided in the study (**Supplementary Table 6**)^7^.

The cell type annotated scRNA expression matrix was then integrated with the scATAC gene score matrix (described in the above section) using the “addGeneIntegrationMatrix” function from ArchR, which identifies corresponding cells across datasets or “anchors” using Seurat’s mutual nearest neighbors (MNN) algorithm. To scale this step across thousands of cells, the total number of cells was divided into smaller groups and alignments were performed in parallel. Cell type labels within the Seurat scRNA-seq object metadata were transferred to the corresponding mutual nearest neighbors in the scATAC-seq data along with their gene expression signature. The output of the integration step resulted in scATAC-seq cells having both a chromatin accessibility and gene expression profile. After integration, scATAC cells were re-annotated in UMAP space using the scRNA transferred labels and these newly defined groups were used for downstream analyses as an alternative annotation in addition to the marker gene-based annotation. The scRNA transferred labels were also used in the Fibromyocyte vs. SMC differential analysis (**Figure 3h-j**).

### Cell-type specific peaks and transcription factor motif enrichment

Genome-wide chromatin accessible regions for each “pseudo bulk” sample (reads from the same cluster were combined as a new sample) were detected using the “addReproduciblePeakSet” function in ArchR (with parameters extsize=100, cutOff=0.01, extendSummits=200). 323,767 chromatin accessible regions (peaks) were detected thereafter. The cell-type specific peaks (marker peaks) for each cluster/cell-type were identified using a similar strategy as identification of cell-type specific genes (with parameters FDR <= 0.01 & Log2FC >= 1). This resulted in a total of 173,357 cell-type specific peaks for different cell types. The enriched motifs for each cell type were predicted using the “addMotifAnnotations” function in the ArchR package based on the HOMER^42^ motif database (v4.11). The chromatin accessibility variability and deviation of transcription factors was estimated by chromVAR (1.12.0)^57^ with genome-wide motif sites provided as potential binding sites.

### Peak pathway annotation

To perform functional annotation of cell-type marker peaks (**Supplementary Figure 6**), we used GREAT^95^ with default parameters. The top 5 function terms (in Biological Processes) for each cell type were displayed as a dot plot. The colors and sizes of the dots represent −log10(FDR) (from the hypergeometric gene-based test) and the percentage of associated gene, respectively.

### Trajectory analysis

The trajectory analysis was performed using the “addTrajectory” function in ArchR and specifying the cluster order (cluster 6 – cluster 5 – cluster 7, Figure 3f). We further visualized trajectory-dependent changes of (summarized) ATAC-seq motif signals using “plotTrajectoryHeatmap” function in ArchR (Figure 3g).

### Chromatin accessible peak to gene linkage and co-accessibility analysis

We leveraged the integrated scRNA and scATAC data in order to explore correlations between co-accessible regions and gene expression. These candidate gene regulatory interactions were predicted using the “getPeak2GeneLinks” function with default parameters in ArchR. The peak2gene loops were collected and plotted using the Sushi package^96^ with red color highlighting SNP associated loops and grey color for other loops (**Figure 6a-b**). For the loops around *VEGFA* and *CDH13* promoter (**Figure S6a-b**), the loops were predicted using the “addCoAccessibility” function with the additional parameter “maxDist=1e6”.

### Differential analysis between SMCs and fibromyocytes

#### Data processing

In order to explore differential regulatory profiles in SMCs and fibromyocytes, we performed another round of scATAC-scRNA-seq integration (using the data from Wirka et al. 2019). To delineate modulated SMCs (fibromyocytes) with high resolution and to further validate results from Wirka et al, we slightly increased the stringency of quality control for scRNA-seq data including coverage as an additional metric. Genes expressed in less than 5 cells were filtered out. Cells expressing < 500 and > 2500 genes, and with < 2000 UMIs were also trimmed from the dataset to prune defective cells, multiplets or cells with low coverage. Moreover, cells containing <1% and > 5% of reads mapping to the mitochondrial genome were discarded. Upon discarding lower quality cells, 7209 high quality cells remained for subsequent analysis. Read counts normalization and dimensionality reduction was performed as described above. To optimize clustering, several resolutions (granularity parameters) were applied to avoid under- or over-clustering of the data. A resolution of 1.2-1.7 yielded a consistent number of clusters and upon identifying the differentially expressed genes for each, clusters could be successfully annotated using the human scRNA cluster-specific gene lists provided by Wirka et al 2019.

#### Differential accessibility, annotation and promoter analysis

Using cell type groupings defined by scRNA-seq label transfer, differentially expressed peaks between traditional Smooth Muscle Cells (SMC) and Fibromyocytes were identified using a Wilcoxon-test as implemented in the ArchR package. Peaks called during this analysis had a width of 500bp. The threshold for differential peak significance was set at FDR <= 0.05 and Log2 fold change > 1, resulting in a total of 5281 significantly upregulated peaks and 2121 downregulated peaks. For differentially accessible promoter analysis, the promoter coordinates of protein coding genes were extracted using the R packages ensembldb v2.14.0^97^ and EnsDb.Hsapiens.v86 v2.99.0 (Rainer J (2017). *EnsDb.Hsapiens.v86: Ensembl based annotation package*. R package version 2.99.0). These promoter coordinates were overlapped with upregulated and downregulated peak coordinates using the R package GenomicRanges v1.42.0^98^.

As an additional approach for differential peaks annotation, protein coding genes coordinates were extracted with ensembldb. Upregulated and downregulated peaks were annotated with the nearest protein coding gene using GenomicRanges v1.42.0. This annotation was validated using the R package ChIPseeker v1.26.0^99^ along with TxDb.Hsapiens.UCSC.hg38.knownGene v3.10.0 (Team BC, Maintainer BP (2019). *TxDb.Hsapiens.UCSC.hg38.knownGene: Annotation package for TxDb object(s)*. R package version 3.4.6.).

#### Differential motif and GO enrichment

Differential upregulated and downregulated peaks for fibromyocytes were converted into BED files and tested for TF motif enrichment using the command line tool HOMER v4.10 (http://homer.ucsd.edu/homer/ngs/). The findMotifsGenome.pl script was used to search for known and *de-novo* motifs. The analysis was run using default values, with the exception of the parameter “-size”, that was set to “-size given” to define peaks width from the BED files data instead of arbitrarily defining a constant value. The same BED files were used for region set enrichment analysis using the Genomic Regions Enrichment of Annotations Tool (GREAT v4.0.4) (http://great.stanford.edu/public/html/index.php) using the whole genome as background (reported results are from the GO database).

### LD score regression

We used the LDSC package (https://github.com/bulik/ldsc) to perform LD Score Regression using our single-cell ATAC peaks^59^. We first downloaded GWAS summary statistics for: CAD^13^; carotid intima-media thickness (cIMT)^100^; carotid artery plaque^100^; diastolic blood pressure (DBP), systolic blood pressure (SBP), and pulse pressure (PP) from the Million Veterans Program (MVP)^101^; Alzheimer’s Disease^102^; type 2 diabetes (UK Biobank)^103^; body mass index (BMI) (UK Biobank)^103^; and White Blood Cell (WBC) count (UK Biobank)^103^. The UK Biobank summary statistics were downloaded from https://alkesgroup.broadinstitute.org/UKBB/. We used the provided munge_sumstats.py script to convert these GWAS summary statistics to a format compatible with ldsc. For each coronary artery cell type we lifted over bed file peak coordinates from hg38 to hg19. We then used these hg19 bed files to make annotation files for each cell type. We performed LD Score Regression according to the cell type-specific analysis tutorial (https://github.com/bulik/ldsc/wiki/Cell-type-specific-analyses).

### CAD GWAS datasets

For comparison with CAD GWAS data we primarily used summary statistics from van der Harst et al.^13^ that performed GWAS in UK Biobank subjects followed by replication in CARDIoGRAMplusC4D. For overlap of cell type peaks we also used this SNPs from this GWAS and a recent CAD GWAS (Koyama et al. Nature Genetics 2021) that performed *trans*-ancestry meta-analysis^15^. We obtained the list of lead SNPs (p < 5 x 10^-8^) from GWAS catalog (https://www.ebi.ac.uk/gwas) for the van der Harst study and Supplementary Table 8 of Koyama et al. We used these lead variants as input to LDlinkR (https://ldlink.nci.nih.gov, https://github.com/CBIIT/LDlinkR) and subsequently kept variants with r^2^ >= 0.8 using EUR population.

### Overlap of CAD GWAS SNPs with cell type peaks

We first used the UCSC liftover tool to convert the LDlinkR output of LD-expanded CAD SNPs from hg19 to hg38 coordinates. To account for CAD variants directly adjacent to a peak we considered a 100 bp window with the SNP lying within the center. We used BEDOPS^104^ to extend the SNP position 50 bp in each direction. Next we used bedtools intersect^105^ to intersect these SNP coordinates with peak bed files from each cell type. We considered 1) peaks from the total peak set annotated to that cell type and 2) cell type marker peaks.

### Genomic DNA sequencing

We isolated genomic DNA for each patient using the Qiagen DNeasy Blood and Tissue Kit. Approximately 20-25 mg of frozen left ventricle or coronary artery was placed in a 1.5 mL microcentrifuge tube and tissue lysed using lysis buffer, proteinase K, heating at 56°C for 1-3 hours or overnight, and intermittent vortexing. We then followed the kit instructions and genomic DNA eluted using 100 μl of Buffer AE (TE buffer). Genomic DNA samples were diluted to concentrations of between 5 ng/μl and 15 ng/μl in skirted 96 well PCR plates using TE buffer. Plates were sealed and shipped to Gencove (New York, USA) for 0.4X low-pass genomic DNA sequencing.

### Genotype phasing, imputation, and liftover of genomic coordinates

We obtained and downloaded the low-pass 0.4x whole-genome sequencing files (unphased) from the Gencove website. These were all provided in human genome build b37. We phased and imputed to the 1000 Genomes reference panel using Beagle (v5.1)^106,107^. We then used Picard to liftover the phased autosomal VCFs from b37 to hg19, then hg19 to hg38 (“Picard Toolkit.” 2019. Broad Institute, GitHub Repository. http://broadinstitute.github.io/picard/; Broad Institute). Approximately 43,000 variants could not be mapped after liftover and were subsequently discarded. This left approximately 38 million total variants (10.1 million variants with minor allele frequency >1 %).

### Chromatin accessibility QTL preprocessing

To identify cell type-specific chromatin accessibility QTLs (caQTLs) we focused on four coronary cell types: smooth muscle cells, endothelial cells, fibroblasts, and macrophages. We first extracted cell-type assigned reads from our ArchR analysis in bam format for each scATAC library. For each individual cell type we excluded individuals with less than 5 cells. We ended up with SMC bam files for 41 patients, endothelial cell bam files for 37 patients, fibroblast bam files for 26 patients (due to some samples lacking adventitia), and macrophage bam files for 41 patients. To obtain region sets we took the peak set across all cell types and converted these peaks from bed to saf format as input for featureCounts. We used these peak coordinates in saf format and cell type bam files as input for featureCounts^108^ with the −p flag for paired-end mode. This subsequently generated raw count matrices for SMC, endothelial, fibroblast, and macrophage cells. For each cell type we only retained peaks with an average of 5 read counts across individuals.

We used RASQUAL^73^ to calculate caQTLs that leverages differences between individuals as well as allele-differences within an individual at heterozygous sites^73^. To simplify preparation of RASQUAL input files we used rasqualTools (https://github.com/kauralasoo/rasqual/tree/master/rasqualTools) to prepare compatible scATAC read count, metadata, and sample specific offset files. To calculate sample offsets we adjusted for library size as well the GC content of each peak.

For each patient we obtained genotypes by performing low-pass whole genome sequencing using Gencove. We used VCFtools^109^ to filter for variants with at least 5% minor allele frequency and select patients with corresponding scATAC-seq libraries. We used RASQUAL to create allele-specific vcf files (createASVCF.sh) for each cell type, which contains genotype information plus counts for reference and alternative alleles. We bypassed the qcFilterBam part of the createASVCF.sh script due to incompatibility and memory issues with our single-cell bam files. However, our bam files extracted from each cell type were previously filtered using ArchR and contain high quality cells and reads.

### Calculation of chromatin accessibility QTLs

For each scATAC peak we tested association for all variants within a 10 kb window. We ran rasqual using the -t flag to output only the top associated SNP for each peak. To obtain a null distribution of q values we performed 5 separate permutation runs for each cell type using the --random-permutation flag to break the relationship between genotype and peak accessibility. To adjust for multiple testing we performed two FDR (false discovery rate) corrections. First, for each peak we obtained a q value corresponding to the SNP level FDR for that peak. https://github.com/natsuhiko/rasqual/issues/21. Next, for each peak we average the q values across the 5 RASQUAL permutation runs. This produced two vectors: one with real RASQUAL q values and one from the permuted q values for each peak. By comparing the real and permuted vectors of q values we were then able to calculate the q value corresponding to either 10% FDR or 5% FDR. For plotting RASQUAL caQTL results, we took raw count files for each cell type, adjusted for library size and performed variance stabilizing transformation in DESeq2^110^.

### Overlap of caQTLs with GTEx eQTLs

We used the QTlizer R package^75^ to query the significant SMC caQTL rsIDs for eQTL signals in GTEx v8. We only retained GTEx eQTL signals at 5% FDR and subsequently filtered for relevant arterial tissues (coronary artery, aorta, tibial artery).

### Publicly available gene expression data

Gene expression levels, expression quantitative loci (eQTL) data, and eQTL boxplots were obtained from the Genotype-Tissue Expression (GTEx) v8 portal website (https://www.gtexportal.org/home/). Differential gene expression data from publicly available Gene Expression Omnibus (GEO) in cardiovascular relevant systems was obtained via the HeartBioPortal (https://www.heartbioportal.com).

### Functional variant predictive modeling based on accessible sequences

We first downloaded CAD GWAS summary statistics from van der Harst et al.^13^ and retained variants passing the genome-wide threshold (p < 5 x 10^-8^). This resulted in 10,117 variants that were tested. The variant scoring analysis (**Figure 5h**) was conducted using the lsgkm package (https://github.com/kundajelab/lsgkm)^76,111^ and the GkmExplain package (https://github.com/kundajelab/gkmexplain)^77^. For a given cell type (e.g. SMC), the reads from all the individual cells assigned to the cell type were first collected as a pseudo bulk sample. The pseudo bulk ATAC-seq peaks were detected with MACS2^112^ (paired end mode, with additional parameter -q 0.01). In the model building step, peaks were split 10-fold for crossvalidation. For each fold, the top 60000 peaks with highest −log10(q-value) were selected as the training set. The +/− 500 bp sequence from the peak summits were used as a positive set, while sequences form a 1000 bp region outside of peaks with matching GC-content were used as a negative set. The importance score of all the positions around the target SNP (up to +/− 100 bp) were plotted as sequence logo (**Figure 5h**).

### STARNET gene regulatory network analysis and module enrichment

Based on STARNET multi-tissue gene expression data (bulk RNA-seq), tissue-specific and cross-tissue co-expression modules were inferred using WGCNA^113^ as previously described^114^. Enrichment for clinical trait associations was computed by aggregating Pearson’s correlation p values by co-expression module using Fisher’s method. Enrichment for differentially expressed genes was calculated using the hypergeometric test, with differentially expressed genes called by DESeq2 (+/− 30% change, FDR<0.01) with adjustment for age and gender. The gene regulatory network was inferred among PRDM16 and TBX2 co-expressed genes using GENIE3^115^ with potential regulators restricted to eQTL genes or known transcription factors. To identify hub genes in the network, weighted key driver analysis (wKDA) was carried out using the Mergeomics R package^116^.

### Immunofluorescence of human coronary artery tissues

Human coronary artery tissues were obtained as described above. Briefly, coronary artery segments were isolated from healthy and subclinical atherosclerotic left main and right coronary artery branches. Tissues were embedded in OCT blocks, snap-frozen in liquid nitrogen and stored at −80° C. Tissue blocks were cryosectioned at −20°C at 6 μm thickness and processed for immunostaining. Sections were rehydrated in PBS at room temperature (RT) and fixed in 10% neutral buffered formaldehyde for 10 min at RT, followed by PBS washes, protein blocking in casein buffer for 1 hr at RT, and incubated overnight at 4° C with anti-LMOD1 rabbit polyclonal antibody (Proteintech, 15117-1-AP; 1:100) or anti-PRDM16 rabbit polyclonal antibody (Abcam, ab106410; 1:2000) or anti-rabbit IgG negative control (Thermo Fisher, 02-6102), with optimal dilutions determined by titrations with control tissues. Sections were washed in PBS and incubated with donkey anti-rabbit Alexa Fluor 555 conjugated secondary antibody (Thermo Fisher, A32794; 1:150) for 30 min at RT, washed in PBS, and stained with DAPI (1:500) and coverslipped using aqueous mounting media.

Whole slide images were captured at 25X magnification using a Zeiss LSM 880 Indimo, AxioExaminer confocal microscope with a Plan-Apochromat 25×/0.8 M27 objective in Line Sequential unidirectional mode. Signal corresponding to the DAPI (channel 1) and the protein of interest (channel 2) were obtained using lasers (respective wavelength of excitation of 405 and 561nm); a PMT and filters were used to collect the fluorescence emitted respectively at 410-480nm and 561-597nm. Images in both channels were merged with the Zeiss ZEN 3.3 Lite software (version 3.3.89). Brightness, Gamma, and Contrast were uniformly adjusted. Corresponding regions of interest of the sections immunostained with both antibodies were numerically magnified. Whole slide images were reconstructed from tiles acquired in brightfield using a high resolution HV-F203SCL Hitachi camera mounted on a Axio Scan microscope using a Plan-Apochromat 10X/0.3 objective.

For histology analysis, sections were stained with hematoxylin and eosin. Images were captured using a Zeiss 183 Axio Scan Z1 at 20X magnification. The resulting czi files were visualized for staining using Zeiss ZEN 3.3 Lite software (version 3.3.89).

### Genome browser tracks

For each cell type we created bigWig files from aggregated cells and created custom tracks that were uploaded to the UCSC Genome Browser and viewed using hg38.

## Data availability

All raw and processed single-cell chromatin accessibility sequencing datasets are made available on the Gene Expression Omnibus (GEO) database (Accession GSE175621).

## Code availability

Our results make use of published software tools with detailed parameters included in the Methods. All custom scripts used to generate these results are available on GitHub (https://github.com/MillerLab-CPHG/Coronary_scATAC).

## Acknowledgements

This work was supported by grants from: the National Institutes of Health (R01HL148239 and R00HL125912 to C.L.M.; R35GM133712 to C.Z.; R01HL141425 to A.V.F; R01HL125863 to J.LM.B.; R01HL130423, R01HL135093, and R01HL148167-01A1 to J.C.K.; R35HL144475, R01 HL125224, R01HL134817 and R01HL139478 to T.Q. and R01HL123370 to N.J.L.), the American Heart Association (20POST35120545 to A.W.T; A14SFRN20840000 to J.LM.B; 19EIA34770065 to N.J.L.), the Swedish Research Council and Heart Lung Foundation (2018-02529 and 20170265 to J.LM.B.), and the Fondation Leducq (‘PlaqOmics’ 18CVD02 to N.J.L., J.LM.B., A.V.F. and C.L.M.). We thank Peter Chiu, Paul Chang and Michael Wong for surgical assistance and research donor heart procurement. We thank Tiffany Koyano for assistance in extracting clinical information. We thank all of the transplant recipient and heart donors, family members, study coordinators and transplant tissue procurement team at Stanford. We thank Boxiang Liu and Natsuhiko Kumasaka for helpful discussions on QTL scripts. Finally, we thank all staff of the genome core facilities for library construction and sequencing assistance.

## Author Contributions

C.L.M, and C.Z., jointly supervised research primarily related to the study. J.LM.B., J.C.K., N.J.L., A.V.F., and T.Q. jointly supervised research secondarily related to the study. A.W.T., S.H., C.Z. and C.L.M. conceived and designed the experiments. A.W.T., K.S-C., E.A.F. and S.K.B.G. performed the experiments. A.W.T., S.H., J.V.M. performed the statistical analyses. A.W.T., S.H., J.V.M. W.F.M., C.J.H., D.W., G.A., and C.L.M. analyzed the data. K.S-C., E.A.F., S.K., A.B.F., N.G.L., L.M., S.K.B.G., S.O-G., E.A.A., T.Q., A.V.F., N.J.L., J.C.K. and J.LM.B. contributed reagents/materials/analysis tools. A.W.T., S.H., J.V.M. W.F.M., C.J.H., D.W., G.A., C.Z. and C.L.M. wrote the paper.

## Competing Interests

Dr. Björkegren is a shareholder in Clinical Gene Network AB who have an invested interest in STARNET. Dr. Finn at CVPath also acknowledges receiving financial support from the following entities: 4C Medical, 4Tech, Abbott Vascular, Ablative Solutions, Absorption Systems, Advanced NanoTherapies, Aerwave Medical, Alivas, Amgen, Asahi Medical, Aurios Medical, Avantec Vascular, BD, Biosensors, Biotronik, Biotyx Medical, Bolt Medical, Boston Scientific, Canon, Cardiac Implants, Cardiawave, CardioMech, Cardionomic, Celonova, Cerus, EndoVascular, Chansu Vascular Technologies, Childrens National, Concept Medical, Cook Medical, Cooper Health, Cormaze, CRL, Croivalve, CSI, Dexcom, Edwards Lifesciences, Elucid Bioimaging, eLum Technologies, Emboline, Endotronix, Envision, Filterlex, Imperative Care, Innovalve, Innovative, Cardiovascular Solutions, Intact Vascular, Interface Biolgics, Intershunt Technologies, Invatin, Lahav, Limflow, L&J Bio, Lutonix, Lyra Therapeutics, Mayo Clinic, Maywell, MDS, MedAlliance, Medanex, Medtronic, Mercator, Microport, Microvention, Neovasc, Nephronyx, Nova Vascular, Nyra Medical, Occultech, Olympus, Ohio Health, OrbusNeich, Ossio, Phenox, Pi-Cardia, Polares Medical, Polyvascular, Profusa, ProKidney, LLC, Protembis, Pulse Biosciences, Qool Therapeutics, Recombinetics, Recor Medical, Regencor, Renata Medical, Restore Medical, Ripple Therapeutics, Rush University, Sanofi, Shockwave, SMT, SoundPipe, Spartan Micro, Spectrawave, Surmodics, Terumo Corporation, The Jacobs Institute, Transmural Systems, Transverse Medical, TruLeaf, UCSF, UPMC, Vascudyne, Vesper, Vetex Medical, Whiteswell, WL Gore, Xeltis. All other authors declare that they have no competing interests relevant to the contents of this paper to disclose.

## Notes

### Competing Interest Statement

Dr. Bjorkegren is a shareholder in Clinical Gene Network AB who have an invested interest in STARNET. All other authors declare that they have no competing interests relevant to the contents of this paper to disclose.

## References

1. Libby, P. Inflammation in atherosclerosis. Nature 420, 868–874 (2002).

2. Souilhol, C., Harmsen, M. C., Evans, P. C. & Krenning, G. Endothelial-mesenchymal transition in atherosclerosis. Cardiovasc. Res. 114, 565–577 (2018).

3. Stary, H. C. et al. A definition of advanced types of atherosclerotic lesions and a histological classification of atherosclerosis. A report from the Committee on Vascular Lesions of the Council on Arteriosclerosis, American Heart Association. Circulation 92, 1355–1374 (1995).

4. Winkels, H. et al. Atlas of the Immune Cell Repertoire in Mouse Atherosclerosis Defined by Single-Cell RNA-Sequencing and Mass Cytometry. Circ. Res. 122, 1675–1688 (2018).

5. Cochain, C. et al. Single-Cell RNA-Seq Reveals the Transcriptional Landscape and Heterogeneity of Aortic Macrophages in Murine Atherosclerosis. Circ. Res. 122, 1661–1674 (2018).

6. Fernandez, D. M. et al. Single-cell immune landscape of human atherosclerotic plaques. Nat. Med. 25, 1576–1588 (2019).

7. Wirka, R. C. et al. Atheroprotective roles of smooth muscle cell phenotypic modulation and the TCF21 disease gene as revealed by single-cell analysis. Nat. Med. 25, 1280–1289 (2019).

8. Depuydt, M. A. C. et al. Microanatomy of the Human Atherosclerotic Plaque by SingleCell Transcriptomics. Circ. Res. 127, 1437–1455 (2020).

9. Pan, H. et al. Single-Cell Genomics Reveals a Novel Cell State During Smooth Muscle Cell Phenotypic Switching and Potential Therapeutic Targets for Atherosclerosis in Mouse and Human. Circulation 142, 2060–2075 (2020).

10. Alencar, G. F. et al. The Stem Cell Pluripotency Genes Klf4 and Oct4 Regulate Complex SMC Phenotypic Changes Critical in Late-Stage Atherosclerotic Lesion Pathogenesis. Circulation (2020) doi:10.1161/CIRCULATIONAHA.120.046672.

11. Wang, Y. et al. Clonally expanding smooth muscle cells promote atherosclerosis by escaping efferocytosis and activating the complement cascade. Proc Natl Acad Sci USA 117, 15818–15826 (2020).

12. Nikpay, M. et al. A comprehensive 1,000 Genomes-based genome-wide association meta-analysis of coronary artery disease. Nat. Genet. 47, 1121–1130 (2015).

13. van der Harst, P. & Verweij, N. Identification of 64 novel genetic loci provides an expanded view on the genetic architecture of coronary artery disease. Circ. Res. 122, 433–443 (2018).

14. Nelson, C. P. et al. Association analyses based on false discovery rate implicate new loci for coronary artery disease. Nat. Genet. 49, 1385–1391 (2017).

15. Koyama, S. et al. Population-specific and trans-ancestry genome-wide analyses identify distinct and shared genetic risk loci for coronary artery disease. Nat. Genet. 52, 1169–1177 (2020).

16. Erdmann, J., Kessler, T., Munoz Venegas, L. & Schunkert, H. A decade of genome-wide association studies for coronary artery disease: the challenges ahead. Cardiovasc. Res. 114, 1241–1257 (2018).

17. Maurano, M. T. et al. Systematic localization of common disease-associated variation in regulatory DNA. Science 337, 1190–1195 (2012).

18. Edwards, S. L., Beesley, J., French, J. D. & Dunning, A. M. Beyond GWASs: illuminating the dark road from association to function. Am. J. Hum. Genet. 93, 779–797 (2013).

19. Heinz, S., Romanoski, C. E., Benner, C. & Glass, C. K. The selection and function of cell type-specific enhancers. Nat. Rev. Mol. Cell Biol. 16, 144–154 (2015).

20. ENCODE Project Consortium. An integrated encyclopedia of DNA elements in the human genome. Nature 489, 57–74 (2012).

21. Buenrostro, J. D., Giresi, P. G., Zaba, L. C., Chang, H. Y. & Greenleaf, W. J. Transposition of native chromatin for fast and sensitive epigenomic profiling of open chromatin, DNA-binding proteins and nucleosome position. Nat. Methods 10, 1213–1218 (2013).

22. Miller, C. L. et al. Integrative functional genomics identifies regulatory mechanisms at coronary artery disease loci. Nat. Commun. 7, 12092 (2016).

23. Liu, B., Gloudemans, M. J., Rao, A. S., Ingelsson, E. & Montgomery, S. B. Abundant associations with gene expression complicate GWAS follow-up. Nat. Genet. 51, 768–769 (2019).

24. Zhao, Q. et al. Molecular mechanisms of coronary disease revealed using quantitative trait loci for TCF21 binding, chromatin accessibility, and chromosomal looping. Genome Biol. 21, 135 (2020).

25. Stolze, L. K. et al. Systems Genetics in Human Endothelial Cells Identifies Non-coding Variants Modifying Enhancers, Expression, and Complex Disease Traits. Am. J. Hum. Genet. 106, 748–763 (2020).

26. Buenrostro, J. D. et al. Single-cell chromatin accessibility reveals principles of regulatory variation. Nature 523, 486–490 (2015).

27. Cusanovich, D. A. et al. Multiplex single cell profiling of chromatin accessibility by combinatorial cellular indexing. Science 348, 910–914 (2015).

28. Satpathy, A. T. et al. Massively parallel single-cell chromatin landscapes of human immune cell development and intratumoral T cell exhaustion. Nat. Biotechnol. 37, 925–936 (2019).

29. Corces, M. R. et al. Single-cell epigenomic analyses implicate candidate causal variants at inherited risk loci for Alzheimer’s and Parkinson’s diseases. Nat. Genet. 52, 1158–1168 (2020).

30. Chiou, J. et al. Single-cell chromatin accessibility identifies pancreatic islet cell type- and state-specific regulatory programs of diabetes risk. Nat. Genet. 53, 455–466.

31. Chiou, J. et al. Interpreting type 1 diabetes risk with genetics and single-cell epigenomics. Nature (2021) doi:10.1038/s41586-021-03552-w.

32. Hocker, J. D. et al. Cardiac cell type-specific gene regulatory programs and disease risk association. Sci. Adv. 7, (2021).

33. Muto, Y. et al. Single cell transcriptional and chromatin accessibility profiling redefine cellular heterogeneity in the adult human kidney. Nat. Commun. 12, 2190 (2021).

34. Domcke, S. et al. A human cell atlas of fetal chromatin accessibility. Science 370, (2020).

35. Rai, V. et al. Single-cell ATAC-Seq in human pancreatic islets and deep learning upscaling of rare cells reveals cell-specific type 2 diabetes regulatory signatures. Mol. Metab. 32, 109–121 (2020).

36. Corces, M. R. et al. An improved ATAC-seq protocol reduces background and enables interrogation of frozen tissues. Nat. Methods 14, 959–962 (2017).

37. Granja, J. M. et al. ArchR is a scalable software package for integrative single-cell chromatin accessibility analysis. Nat. Genet. 53, 403–411.

38. Granja, J. M. et al. Single-cell multiomic analysis identifies regulatory programs in mixed-phenotype acute leukemia. Nat. Biotechnol. 37, 1458–1465 (2019).

39. Virmani, R., Burke, A. P., Farb, A. & Kolodgie, F. D. Pathology of the vulnerable plaque. J. Am. Coll. Cardiol. 47, C13–8 (2006).

40. Mulligan-Kehoe, M. J. & Simons, M. Vasa vasorum in normal and diseased arteries. Circulation 129, 2557–2566 (2014).

41. Virmani, R. et al. Atherosclerotic plaque progression and vulnerability to rupture: angiogenesis as a source of intraplaque hemorrhage. Arterioscler. Thromb. Vasc. Biol. 25, 2054–2061 (2005).

42. Heinz, S. et al. Simple combinations of lineage-determining transcription factors prime cis-regulatory elements required for macrophage and B cell identities. Mol. Cell 38, 576–589 (2010).

43. Creemers, E. E., Sutherland, L. B., Oh, J., Barbosa, A. C. & Olson, E. N. Coactivation of MEF2 by the SAP domain proteins myocardin and MASTR. Mol. Cell 23, 83–96 (2006).

44. Maeda, T., Gupta, M. P. & Stewart, A. F. R. TEF-1 and MEF2 transcription factors interact to regulate muscle-specific promoters. Biochem. Biophys. Res. Commun. 294, 791–797 (2002).

45. Almontashiri, N. A. M. et al. 9p21.3 Coronary Artery Disease Risk Variants Disrupt TEAD Transcription Factor-Dependent Transforming Growth Factor β Regulation of p16 Expression in Human Aortic Smooth Muscle Cells. Circulation 132, 1969–1978 (2015).

46. Yoshida, T. et al. Myocardin is a key regulator of CArG-dependent transcription of multiple smooth muscle marker genes. Circ. Res. 92, 856–864 (2003).

47. Du, K. L. et al. Myocardin is a critical serum response factor cofactor in the transcriptional program regulating smooth muscle cell differentiation. Mol. Cell. Biol. 23, 2425–2437 (2003).

48. Chen, J., Kitchen, C. M., Streb, J. W. & Miano, J. M. Myocardin: a component of a molecular switch for smooth muscle differentiation. J. Mol. Cell. Cardiol. 34, 1345–1356 (2002).

49. Wang, D.-Z. et al. Potentiation of serum response factor activity by a family of myocardin-related transcription factors. Proc Natl Acad Sci USA 99, 14855–14860 (2002).

50. Meadows, S. M., Myers, C. T. & Krieg, P. A. Regulation of endothelial cell development by ETS transcription factors. Semin. Cell Dev. Biol. 22, 976–984 (2011).

51. Stamatovic, S. M., Keep, R. F., Mostarica-Stojkovic, M. & Andjelkovic, A. V. CCL2 regulates angiogenesis via activation of Ets-1 transcription factor. J. Immunol. 177, 2651–2661 (2006).

52. Zhang, D. E., Hetherington, C. J., Chen, H. M. & Tenen, D. G. The macrophage transcription factor PU.1 directs tissue-specific expression of the macrophage colonystimulating factor receptor. Mol. Cell. Biol. 14, 373–381 (1994).

53. Cui, L. et al. Activation of JUN in fibroblasts promotes pro-fibrotic programme and modulates protective immunity. Nat. Commun. 11, 2795 (2020).

54. Kitoh, A. et al. Indispensable role of the Runx1-Cbfbeta transcription complex for in vivo-suppressive function of FoxP3+ regulatory T cells. Immunity 31, 609–620 (2009).

55. Ono, M. et al. Foxp3 controls regulatory T-cell function by interacting with AML1/Runx1. Nature 446, 685–689 (2007).

56. Masuda, A. et al. Essential role of GATA transcriptional factors in the activation of mast cells. J. Immunol. 178, 360–368 (2007).

57. Schep, A. N., Wu, B., Buenrostro, J. D. & Greenleaf, W. J. chromVAR: inferring transcription-factor-associated accessibility from single-cell epigenomic data. Nat. Methods 14, 975–978 (2017).

58. Nagao, M. et al. Coronary Disease-Associated Gene TCF21 Inhibits Smooth Muscle Cell Differentiation by Blocking the Myocardin-Serum Response Factor Pathway. Circ. Res. 126, 517–529 (2020).

59. Bulik-Sullivan, B. et al. LD Score regression distinguishes confounding from polygenicity in genome-wide association studies. Nat. Genet. 47, 291–295 (2015).

60. Tabas, I. & Lichtman, A. H. Monocyte-Macrophages and T Cells in Atherosclerosis. Immunity 47, 621–634 (2017).

61. Farrugia, A. J. et al. CDC42EP5/BORG3 modulates SEPT9 to promote actomyosin function, migration, and invasion. J. Cell Biol. 219, (2020).

62. Nyati, K. K., Agarwal, R. G., Sharma, P. & Kishimoto, T. Arid5a regulation and the roles of arid5a in the inflammatory response and disease. Front. Immunol. 10, 2790 (2019).

63. Nott, A. et al. Brain cell type-specific enhancer-promoter interactome maps and disease-risk association. Science 366, 1134–1139 (2019).

64. Xu, S. et al. The novel coronary artery disease risk gene JCAD/KIAA1462 promotes endothelial dysfunction and atherosclerosis. Eur. Heart J. 40, 2398–2408 (2019).

65. Beaudoin, M. et al. Myocardial Infarction-Associated SNP at 6p24 Interferes With MEF2 Binding and Associates With PHACTR1 Expression Levels in Human Coronary Arteries. Arterioscler. Thromb. Vasc. Biol. 35, 1472–1479 (2015).

66. Nanda, V. et al. Functional regulatory mechanism of smooth muscle cell-restricted LMOD1 coronary artery disease locus. PLoS Genet. 14, e1007755 (2018).

67. Benaglio, P. et al. Mapping genetic effects on cell type-specific chromatin accessibility and annotating complex trait variants using single nucleus ATAC-seq. BioRxiv (2020) doi:10.1101/2020.12.03.387894.

68. Calderon, D. et al. Landscape of stimulation-responsive chromatin across diverse human immune cells. Nat. Genet. 51, 1494–1505 (2019).

69. Gate, R. E. et al. Genetic determinants of co-accessible chromatin regions in activated T cells across humans. Nat. Genet. 50, 1140–1150 (2018).

70. Bryois, J. et al. Evaluation of chromatin accessibility in prefrontal cortex of individuals with schizophrenia. Nat. Commun. 9, 3121 (2018).

71. Khetan, S. et al. Type 2 Diabetes-Associated Genetic Variants Regulate Chromatin Accessibility in Human Islets. Diabetes 67, 2466–2477 (2018).

72. Currin, K. W. et al. Genetic effects on liver chromatin accessibility identify disease regulatory variants. Am. J. Hum. Genet. (2021) doi:10.1016/j.ajhg.2021.05.001.

73. Kumasaka, N., Knights, A. J. & Gaffney, D. J. Fine-mapping cellular QTLs with RASQUAL and ATAC-seq. Nat. Genet. 48, 206–213 (2016).

74. Liu, B. et al. Genetic regulatory mechanisms of smooth muscle cells map to coronary artery disease risk loci. Am. J. Hum. Genet. 103, 377–388 (2018).

75. Munz, M. et al. Qtlizer: comprehensive QTL annotation of GWAS results. Sci. Rep. 10, 20417 (2020).

76. Ghandi, M., Lee, D., Mohammad-Noori, M. & Beer, M. A. Enhanced regulatory sequence prediction using gapped k-mer features. PLoS Comput. Biol. 10, e1003711 (2014).

77. Shrikumar, A., Prakash, E. & Kundaje, A. GkmExplain: fast and accurate interpretation of nonlinear gapped k-mer SVMs. Bioinformatics 35, i173–i182 (2019).

78. Lee, D. et al. A method to predict the impact of regulatory variants from DNA sequence. Nat. Genet. 47, 955–961 (2015).

79. Nasser, J. et al. Genome-wide enhancer maps link risk variants to disease genes. Nature 593, 238–243 (2021).

80. Higgins, E. M. et al. Elucidation of MRAS-mediated Noonan syndrome with cardiac hypertrophy. JCI Insight 2, e91225 (2017).

81. Seale, P. et al. PRDM16 controls a brown fat/skeletal muscle switch. Nature 454, 961–967 (2008).

82. Seale, P. et al. Transcriptional control of brown fat determination by PRDM16. Cell Metab. 6, 38–54 (2007).

83. Kajimura, S. et al. Initiation of myoblast to brown fat switch by a PRDM16-C/EBP-beta transcriptional complex. Nature 460, 1154–1158 (2009).

84. Liu, D. et al. PRDM16 Upregulation Induced by MicroRNA-448 Inhibition Alleviates Atherosclerosis via the TGF-β Signaling Pathway Inactivation. Front. Physiol. 11, 846 (2020).

85. Warner, D. R. et al. PRDM16/MEL1: a novel Smad binding protein expressed in murine embryonic orofacial tissue. Biochim. Biophys. Acta 1773, 814–820 (2007).

86. Takahata, M. et al. SKI and MEL1 cooperate to inhibit transforming growth factor-beta signal in gastric cancer cells. J. Biol. Chem. 284, 3334–3344 (2009).

87. Craps, S. et al. Prdm16 supports arterial flow recovery by maintaining endothelial function. Circ. Res. (2021) doi:10.1161/CIRCRESAHA.120.318501.

88. Barron, M. R. et al. Serum response factor, an enriched cardiac mesoderm obligatory factor, is a downstream gene target for Tbx genes. J. Biol. Chem. 280, 11816–11828 (2005).

89. Shirai, M., Imanaka-Yoshida, K., Schneider, M. D., Schwartz, R. J. & Morisaki, T. T-box 2, a mediator of Bmp-Smad signaling, induced hyaluronan synthase 2 and Tgfbeta2 expression and endocardial cushion formation. Proc Natl Acad Sci USA 106, 18604–18609 (2009).

90. Hansson, G. K., Jonasson, L., Holm, J. & Claesson-Welsh, L. Class II MHC antigen expression in the atherosclerotic plaque: smooth muscle cells express HLA-DR, HLA-DQ and the invariant gamma chain. Clin. Exp. Immunol. 64, 261–268 (1986).

91. Cao, J. et al. Joint profiling of chromatin accessibility and gene expression in thousands of single cells. Science 361, 1380–1385 (2018).

92. Ma, S. et al. Chromatin Potential Identified by Shared Single-Cell Profiling of RNA and Chromatin. Cell 183, 1103–1116.e20 (2020).

93. Stuart, T. et al. Comprehensive Integration of Single-Cell Data. Cell 177, 1888–1902.e21 (2019).

94. Korsunsky, I. et al. Fast, sensitive and accurate integration of single-cell data with Harmony. Nat. Methods 16, 1289–1296 (2019).

95. McLean, C. Y. et al. GREAT improves functional interpretation of cis-regulatory regions. Nat. Biotechnol. 28, 495–501 (2010).

96. Phanstiel, D. H., Boyle, A. P., Araya, C. L. & Snyder, M. P. Sushi.R: flexible, quantitative and integrative genomic visualizations for publication-quality multi-panel figures. Bioinformatics 30, 2808–2810 (2014).

97. Rainer, J., Gatto, L. & Weichenberger, C. X. ensembldb: an R package to create and use Ensembl-based annotation resources. Bioinformatics 35, 3151–3153 (2019).

98. Lawrence, M. et al. Software for computing and annotating genomic ranges. PLoS Comput. Biol. 9, e1003118 (2013).

99. Yu, G., Wang, L.-G. & He, Q.-Y. ChIPseeker: an R/Bioconductor package for ChIP peak annotation, comparison and visualization. Bioinformatics 31, 2382–2383 (2015).

100. Franceschini, N. et al. GWAS and colocalization analyses implicate carotid intima-media thickness and carotid plaque loci in cardiovascular outcomes. Nat. Commun. 9, 5141 (2018).

101. Giri, A. et al. Trans-ethnic association study of blood pressure determinants in over 750,000 individuals. Nat. Genet. 51, 51–62 (2019).

102. Jansen, I. E. et al. Genome-wide meta-analysis identifies new loci and functional pathways influencing Alzheimer’s disease risk. Nat. Genet. 51, 404–413 (2019).

103. Sudlow, C. et al. UK biobank: an open access resource for identifying the causes of a wide range of complex diseases of middle and old age. PLoS Med. 12, e1001779 (2015).

104. Neph, S. et al. BEDOPS: high-performance genomic feature operations. Bioinformatics 28, 1919–1920 (2012).

105. Quinlan, A. R. & Hall, I. M. BEDTools: a flexible suite of utilities for comparing genomic features. Bioinformatics 26, 841–842 (2010).

106. Browning, S. R. & Browning, B. L. Rapid and accurate haplotype phasing and missingdata inference for whole-genome association studies by use of localized haplotype clustering. Am. J. Hum. Genet. 81, 1084–1097 (2007).

107. Browning, B. L., Zhou, Y. & Browning, S. R. A One-Penny Imputed Genome from Next-Generation Reference Panels. Am. J. Hum. Genet. 103, 338–348 (2018).

108. Liao, Y., Smyth, G. K. & Shi, W. featureCounts: an efficient general purpose program for assigning sequence reads to genomic features. Bioinformatics 30, 923–930 (2014).

109. Danecek, P. et al. The variant call format and VCFtools. Bioinformatics 27, 2156–2158 (2011).

110. Love, M. I., Huber, W. & Anders, S. Moderated estimation of fold change and dispersion for RNA-seq data with DESeq2. Genome Biol. 15, 550 (2014).

111. Lee, D. LS-GKM: a new gkm-SVM for large-scale datasets. Bioinformatics 32, 2196–2198 (2016).

112. Zhang, Y. et al. Model-based analysis of ChIP-Seq (MACS). Genome Biol. 9, R137 (2008).

113. Langfelder, P. & Horvath, S. WGCNA: an R package for weighted correlation network analysis. BMC Bioinformatics 9, 559 (2008).

114. Talukdar, H. A. et al. Cross-Tissue Regulatory Gene Networks in Coronary Artery Disease. Cell Syst. 2, 196–208 (2016).

115. Huynh-Thu, V. A., Irrthum, A., Wehenkel, L. & Geurts, P. Inferring regulatory networks from expression data using tree-based methods. PLoS ONE 5, (2010).

116. Shu, L. et al. Mergeomics: multidimensional data integration to identify pathogenic perturbations to biological systems. BMC Genomics 17, 874 (2016).

